# CROP: A feature-independent context-aware method for CRISPR-Cas9 frameshift prediction

**DOI:** 10.64898/2026.01.20.700643

**Authors:** Ido Tziony, Yaron Orenstein

**Affiliations:** Department of Computer Science, Bar-Ilan University, Ramat Gan, 5290002, Israel; The Mina and Everard Goodman Faculty of Life Sciences, Bar-Ilan University, Ramat Gan, 5290002, Israel

## Abstract

**Motivation:** The CRISPR-Cas9 complex has revolutionized genome-editing technologies. By designing a 20 nt-long guide RNA, a Cas9 nuclease can be guided to cleave almost any genomic target site (followed by NGG). The cleavage induces double-stranded DNA breaks, which are then repaired by cellular pathways. Accurate CRISPR-Cas9 repair-outcome prediction is essential for designing guide RNAs with desired genomic effects, such as gene knockout. A central challenge is quantifying the rate of frameshifts, i.e. repair-outcomes that lead to a change in the local length that is not a multiplicity of 3. Previous methods for frameshift-rate prediction were trained on only few experimental or cellular contexts, mostly rely on manually defined microhomology features, and are limited by sparse features and class labels.

**Results:** We developed CROP, the first feature-independent context-aware repair-outcome prediction method. By aggregating specific repair outcomes as Δlength classes, CROP overcomes class sparsity. We designed CROP to work with variable input sequence lengths and output classes in order to utilize multiple datasets simultaneously. We benchmarked CROP against state-of-the-art repair-outcome prediction methods over 18 datasets, which we curated and standardized from various studies. Across all datasets, CROP outperformed all competing methods in frameshift-rate prediction. We performed cross-experiment and cross-cellular frameshift predictions to investigate the generalizability of repair mechanisms. Finally, we show that CROP learned microhomology principles from raw sequences without explicit feature engineering, establishing the first end-to-end architecture for CRISPR-Cas9 repair-outcome prediction which learns from multiple datasets.

**Availability and implementation:** CROP is available at https://github.com/OrensteinLab/CROP.

## 1. Introduction

The CRISPR-Cas9 complex [Ishino et al., 1987] is a genomeediting technology adapted from a prokaryotic immune defense mechanism. It uses a programmable 20 nt-long guide RNA to guide a Cas9 nuclease to a genomic *target site*, followed by a 3 bp protospacer adjacent motif (PAM) NGG. The Cas9 nuclease typically induces a double-strand break (DSB) 3 bp upstream of the PAM site, termed the *canonical cut site*, but may also induce a staggered cut, which is shifted on the strand opposite to the PAM [Zuo and Liu, 2016, Shi et al., 2019]. The induced DSB is repaired by cellular repair pathways, which determine the editing outcome and depend on the *target sequence* (i.e., the target site and its flanking sequences). CRISPR-Cas9 genome editing has become a fundamental tool in both research and therapy due to its simplicity, adaptability, and precision [Ishibashi et al., 2020, Dalvie et al., 2022]. However, several computational challenges remain [Liu et al., 2021]: (a) accurately predicting *on-target* efficiency [Elkayam et al., 2024], (b) minimizing *off-target* activity [Yaish and Orenstein, 2024], and (c) accurately predicting the *repair outcomes* [Seale and Gonçalves, 2025].

The repair outcome is governed by the specific pathway utilized to repair the DSB. The three primary repair pathways are non-homologous end joining (NHEJ), microhomology-mediated end joining (MMEJ) (which are both template-free), and homology-directed repair (HDR), which requires a donor template [Yeh et al., 2019]. Since HDR is inefficient, can only occur in the G2 and S phases of the cell, and often generates unwanted byproducts, template-free editing is often preferable in gene-editing applications. The NHEJ pathway typically results in a stochastic distribution of outcomes, characterized by small random insertions or deletions, termed indels. In contrast, MMEJ utilizes microhomology (MH), which is characterized by short, repetitive sequences flanking the DSB. During the MMEJ repair process, these repetitive sequences align, resulting in the deletion of the intervening sequence (Figure 1). The efficiency of the MMEJ process is determined by the biophysical properties of the target sequence, specifically the MH length (i.e., the length of the repetition), and the distances from the cut site to the start of the repetitive sequence on either side (upstream and downstream gaps).

**Fig. 1.**
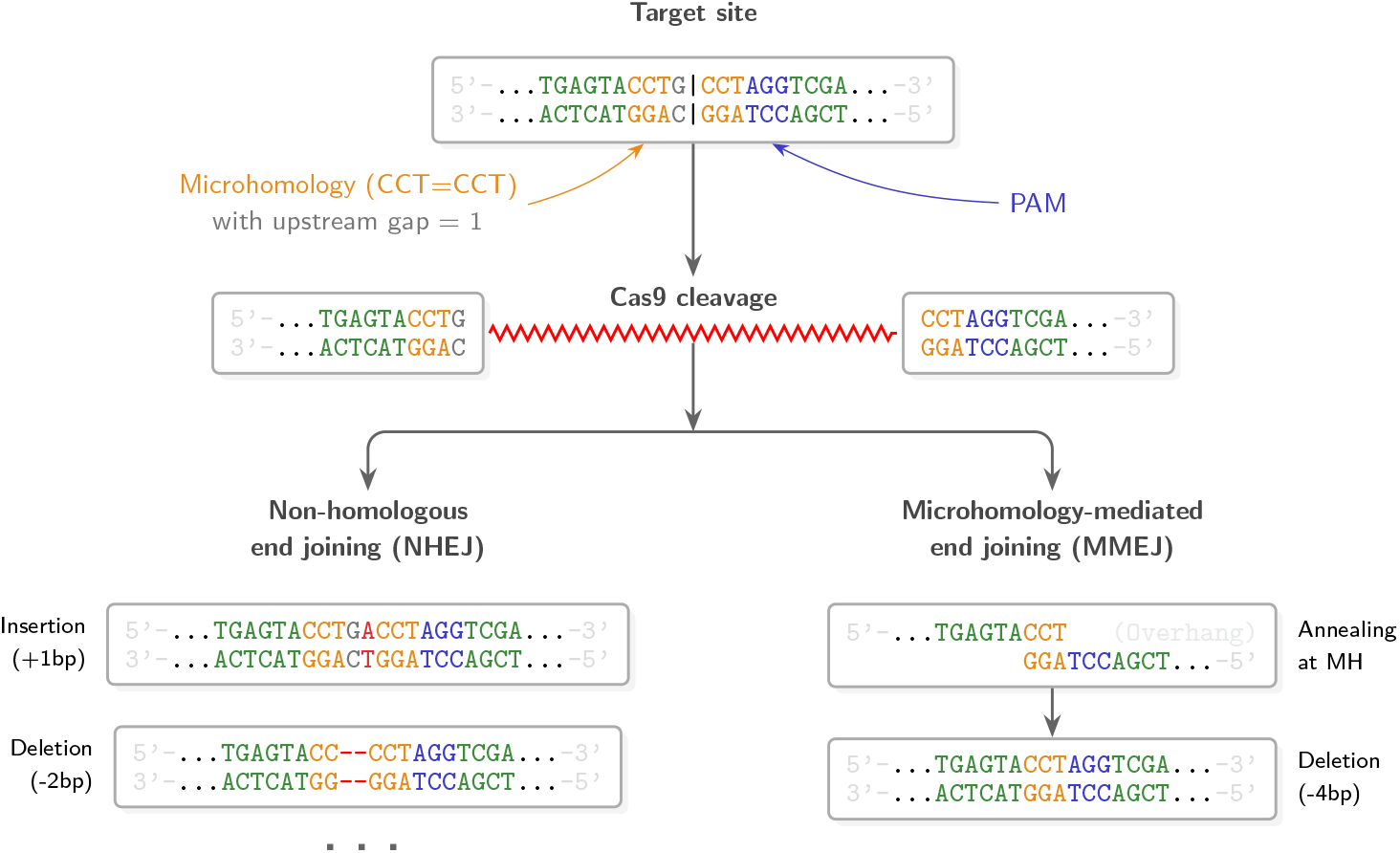
Unified illustrative schema of the two primary template-free cellular repair pathways. Following a double-strand break, repair predominantly proceeds via non-homologous end joining, resulting in variable indels such as insertions or small random deletions, or via microhomology mediated end joining, where annealing of repetitive sequences leads to a deletion of total length of gap size plus repetitive sequence length.

Many repair outcomes result in a *frameshift*, occurring when the indel length is not a multiplicity of 3. Such indels shift the open reading frame, introducing with high likelihood premature stop codons that yield truncated, non-functional proteins. Consequently, gene-editing that results in a frameshift is a popular mechanism for gene knockout [Shalem et al., 2014, Dalvie et al., 2022]. However, repair profiles vary considerably across sequence and cellular contexts, and not all indels shift the reading frame, resulting in variable frameshift rates (FSRs). While various experimental assays were developed and applied to measure repair outcomes of thousands of guide RNAs simultaneously [Zhang et al., 2023], it is infeasible to measure repair outcomes over all possible guide RNAs and over all cellular context. Thus, computational methods were developed to predict repair outcomes based on the target sequence.

Existing methods to repair-outcome prediction typically predict high-resolution, discrete events, specifying both the position and length of an indel [Allen et al., 2019, Shen et al., 2018]. This paradigm has several limitations. First, while predicting specific position-length combinations for simple deletions is already challenging, the combinatorial space for complex indels, comprising both insertions and deletions, is larger and very rarely observed in experimental datasets. Second, for applications such as gene knockout, the specific genomic coordinates of an indel are secondary to its impact on the open reading frame. Third, in MMEJ-driven repair, the alignment of identical sequence repeats creates a mapping degeneracy where a single deletion outcome can be attributed to multiple potential genomic coordinates.

While several methods have been developed to predict repair-outcome distributions (Table 1), they suffer from multiple shortcomings. First, many methods rely on precalculated, hand-crafted MH features, precluding an end-to-end learning process. Conversely, featureless methods struggle to capture MH patterns efficiently since they are constrained by insufficient training data. Furthermore, MH features are inherently sparse and often represented as boolean indicators, which limits effective generalization. Second, many methods impose rigid constraints on input sequence length, PAM positioning, and output classes, rendering them incapable of integrating data from multiple heterogeneous experiments. Third, current methods typically utilize only one dataset for training, with only X-CRISP relying on a second experimental source for fine-tuning, and no method uses multi-tasking. This restricted data exposure limits generalization, as the methods become heavily reliant on cell- or experiment-specific biases. Fourth, some methods cannot predict outcomes which are useful for gene-knockout applications (e.g., SPROUT only reports summary statistics). Last, many methods are no longer maintained or suffer from execution failures, making it impossible to run them.

**Table 1.** State-of-the-art CRISPR repair-outcome prediction methods.

| Method | Architecture | #Samples<sup>a</sup> | #MH features | Max. datasets | Available |
| --- | --- | --- | --- | --- | --- |
| ForeCAST [Allen et al., 2019] | MLR | ≤41.6K | 21 | 1 | ✓ |
| inDelphi [Shen et al., 2018] | NN + KNN | 2.0K | None | 1 | ✓ |
| SPROUT [Leenay et al., 2019] | GB | 1.7K | None | 1 | ✓ |
| Lindel [Chen et al., 2019] | MLR + LR | 4.8K | 2,649 | 1 | ✓ |
| CROTON [Li et al., 2021] | CNN | 35.1K | None | 1 | ✗(Broken) |
| Apindel [Liu et al., 2022] | BiLSTM + Attention | 35.1K | None | 1 | ✗(Broken) |
| Aldit [Zhang et al., 2023] | GB + LR | 80.0K | 2,390 | 1 | ✗(Private) |
| X-CRISP [Seale and Gonçalves, 2025] | NN + MLR | 12.8K | Variable<sup>b</sup> | 2 | ✗(Broken) |
| **CROP (this study)** | **BiGRU + Transformer** | **507.0K** | **None** | Unlimited | ✓ |

<sup>a</sup> Includes the full size (with test set) of each dataset used for training, attributed to the model exposed to the most samples per method.
<sup>b</sup> X-CRISP’s source code suggests: Variable = $5 \times (\text{total possible deletions})$ .
MH: Microhomology; LR: Logistic Regression; MLR: Multinomial Logistic Regression; GB: Gradient Boosting; KNN: K-Nearest Neighbors.

In this study, we present CROP, a feature-independent context-aware method for CRISPR-Cas9 repair-outcome prediction. We curated and standardized 18 datasets from multiple studies for large-scale training and evaluation. CROP’s architecture can integrate heterogeneous datasets of variable input lengths and output classes, and model MH without manual feature engineering. Our comprehensive benchmarking demonstrates that CROP outperforms current state-of-the-art methods in predicting FSR across all evaluated datasets. Furthermore, our ablation analysis underscores the contribution of CROP’s architectural components to its predictive performance. Finally, we employ a multi-tier interpretability analysis to validate that CROP captures MH-based mechanisms. Together, we demonstrate that CROP recovers biological repair logic from target sequences, providing the first end-to-end, interpretable method for context-aware repair-outcome prediction.

## 2. Methods

### 2.1. Dataset curation and standardization

To train CROP, we curated 18 CRISPR-Cas9 repair datasets spanning four studies (Table 2) as follows. We sourced datasets from FORECasT, encompassing multiple cell lineages (K562, HAP1, BOB, CHO, RPE1, and mESC) and alternative Cas9 proteins: eCas9, TREX2, and 2A TREX2. We further incorporated a T-cell dataset from SPROUT and K562, HAP1, and Jurkat datasets from Aldit. Moreover, the Aldit study includes terminal deoxynucleotidyl transferase (DNTT) perturbation datasets (DNTT-OE and DNTT-KO), enabling CROP to possibly capture repair dynamics driven by DNTT [Alt and Baltimore, 1982, Landau et al., 1987].

**Table 2.** Summary of datasets curated and standardized for training and testing in this study.

| Study | Dataset | # TSs | TS length | PAM position | Δlength range |
| --- | --- | --- | --- | --- | --- |
| FORECasT<br>[Allen et al., 2019] | K562 | 35,129 | 79 | {30, 42, 56} | [−70, +75] |
|  | K562 eCas9 | 34,536 |  |  | [−69, +65] |
|  | K562 TREX2 | 35,129 |  |  | [−70, +64] |
|  | K562 2A TREX2 | 35,129 |  |  | [−70, +73] |
|  | HAP1 | 35,128 |  |  | [−70, +73] |
|  | BOB | 35,066 |  |  | [−69, +74] |
|  | CHO | 35,129 |  |  | [−70, +73] |
|  | RPE1 | 34,677 |  |  | [−70, +63] |
|  | mESC | 35,104 |  |  | [−70, +72] |
| SPROUT<br>[Leenay et al., 2019] | Primary T (Normal) | 1,598 | 60 | 33 | [−99, +88] |
|  | Primary T (CA) | 1,598 |  |  | [−99, +20] |
| Aldit<br>[Zhang et al., 2023] | K562 | 79,964 | 85 | 40 | [−29, + 2] |
|  | K562 DNTT-OE | 14,076 |  |  | [−50, +42] |
|  | HAP1 | 15,212 |  |  | [−29, + 2] |
|  | Jurkat | 65,050 |  |  | [−29, + 8] |
|  | Jurkat DNTT-KO | 10,188 |  |  | [−50, +43] |
| inDelphi<br>[Shen et al., 2018] | mESC | 1,962 | 55 | 30 | [−44, +22] |
|  | mESC NHEJ-deficient | 1,961 |  |  | [−44, +16] |
|  | U2OS | 1,950 |  |  | [−44, +20] |

TS: Target Sequence.

For the inDelphi datasets, we utilized the preprocessing provided in the X-CRISP study [Seale and Gonçalves, 2025] for mESC and U2OS lines. This includes a NHEJ-deficient dataset (*Lig*4^−*/*−^ and *Prkdc*^−*/*−^ double-knockouts), which highlights MMEJ patterns. We restricted our use of X-CRISP-processed data to the inDelphi datasets, as the FORECasT datasets which X-CRISP provided contained less than third of the target sequences from the original FORECasT datasets. Furthermore, we observed that the repair-outcome profiles in these target sequences were not perfectly correlated with the original datasets (e.g., FSR Pearson correlation of only 0.786 on the mESC dataset, Supplementary Section S1). Finally, while the Lindel dataset [Chen et al., 2019] was considered, its limited sequence context (17 bp upstream and 6 bp downstream of the canonical cut site) is insufficient for modeling MH-mediated deletions. Our curation process is described in full in Supplementary Section S2.

To standardize the datasets format, we implemented the following procedures. First, to reduce class sparsity, we aggregate repair outcomes by their net change in target sequence length (Δlength). For the Aldit datasets, we excluded insertion or deletion classes labeled as *X*+ (i.e., *X* bp or more) and −*X*+, respectively, as these aggregate categories lack the precise length resolution required for CROP. For the SPROUT dataset, we maintained two versions for fair comparison with existing methods. The first version, which we denote by SPROUT^CA^, follows the preprocessing used by CROTON and Apindel by discarding mixed events (i.e., an insertion and a deletion occurring together) and insertions over 20 bp; we use this version exclusively for direct comparisons with CROTON and Apindel. The second version, denoted by SPROUT^NORMAL^, retains the full experimental distribution and is used to train the final CROP model. Finally, for all datasets, we turned the Δlength rates of each target sequence to a distribution by L1-normalization prior to training and testing CROP.

### 2.2. Train-test partitioning

To prevent train-test leakage, we employed a target-sequencebased partitioning. We assigned all samples associated with a specific target sequence (multiple data points share the same sequence) to either the training or the test set, maintaining a 90:10 ratio.

To facilitate a fair comparison with CROTON and Apindel, we replicated the partitioning and random seed (777) used in CROTON’s source code. Specifically, we partitioned the FORECasT K562 target sequences in the same way as CROTON, and merged the train and validation sets. We then partitioned the remaining target sequences in the FORECasT study.

### 2.3. Evaluating repair-outcome prediction methods

We evaluated publicly available state-of-the-art methods that enable to predict FSR or repair-outcome distributions of multiple target sequences together: FORECasT, inDelphi, and Lindel. We ran FORECasT on a Linux server (Intel Xeon Gold 6338 @ 2.00 GHz, 1× Nvidia A100 80GB, 512 GB RAM). We ran inDelphi and Lindel on a Windows 11 machine (AMD Ryzen 7800×3D @ 5.00GHz, 1× Nvidia RTX 4090 24GB, 32GB RAM).

We excluded SPROUT because it reports summary statistics (e.g., fraction of reads with insertions) rather than predicting repair-outcome distributions or FSRs. We were unable to run CROTON and Apindel due to execution errors despite following the instructions on their repositories (Supplementary Section S3). Consequently, we compared against the reported performance in their studies over the FORECasT K562 and SPROUT^CA^ datasets. We excluded Aldit, as it lacks an opensource implementation and relies on a web interface unsuitable for multiple-target prediction; furthermore, the Aldit study does not report performance for FSR prediction. Finally, we excluded X-CRISP due to execution errors (Supplementary Section S3) and because it was trained on dataset variants that do not perfectly correlate with the original FORECasT dataset.

### 2.4. CROP’s architecture

CROP is based on a hybrid neural network that combines a local recurrent module with a global transformer backbone to predict a Δlength distribution (Figure S4). The model receives two inputs: a one-hot target sequence, and a unique dataset identifier.

To enable CROP to learn MH-associated patterns surrounding the cut site, CROP uses a *MH module* (Supplementary Figure S4A). This module splits the sequence at the canonical cut site and reverses the upstream sequence. This allows a shared bidirectional GRU to process both the upstream and downstream flanks of the canonical cut site. The features learned by the MH module are added to the target sequence embedding via a parametrized gate. This gate is initialized to zero, allowing the model to gradually learn how much weight to assign to the MH module.

The main CROP backbone consists of three transformer encoder layers (Supplementary Figure S4B). CROP utilizes rotary positional embeddings (RoPE) [Su et al., 2023] to model the relative distances between nucleotides. To make CROP aware of a specific dataset, a learnable dataset embedding (Supplementary Figure S4C) is integrated into the network. This embedding is added to the latent representation before every multi-head attention and feed-forward block.

Finally, the backbone extracts a single hidden state immediately downstream of the canonical cut site (Figure 2). The hidden state is processed by a classification head comprising layer normalization, a 256-unit dense layer with GELU activation, and dropout, followed by a linear-softmax layer to predict the Δlength probability distribution.

**Fig. 2.**
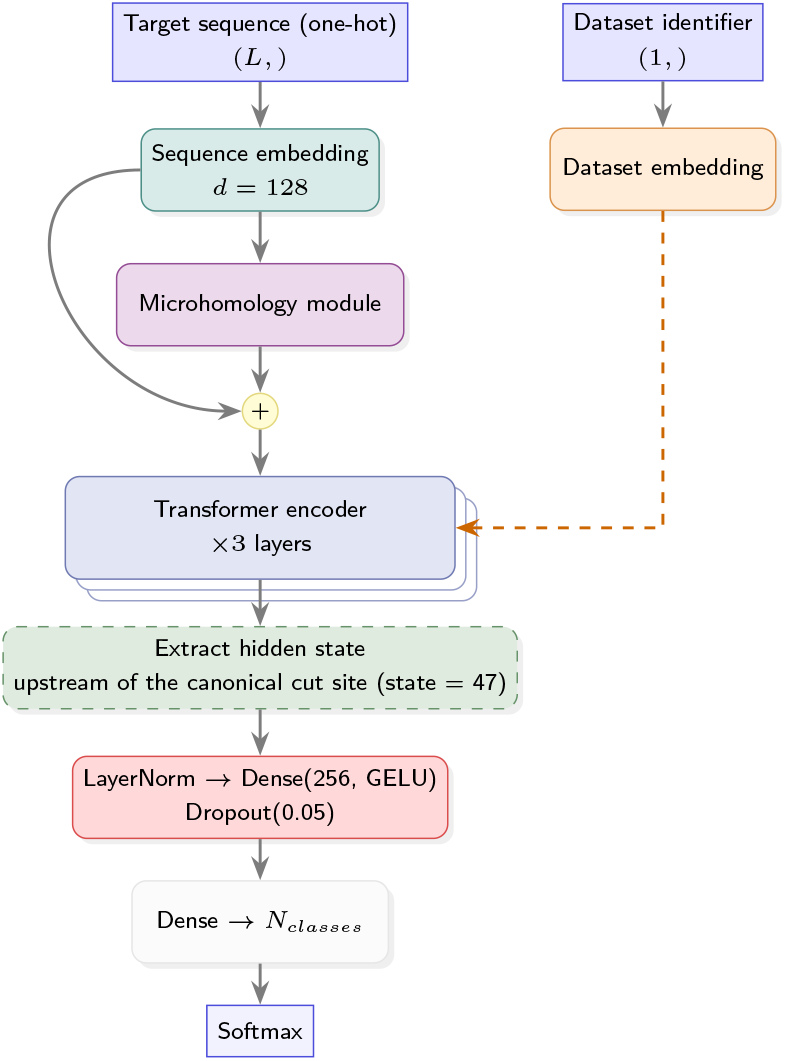
CROP’s architecture

### 2.5. Handling variable input lengths

To ensure fixed input sequence lengths, we standardize all target sequences to a fixed-length window centered on the PAM. We define a window of 103 bp, comprising the 3-bp PAM, 50 bp of upstream flanking sequence, and 50 bp of downstream flanking sequence. Target sequences are then padded with a padding symbol to fit into the fixed input length. During both training and inference, a binary padding mask is generated to ensure that the attention mechanism ignores these padding symbols, preventing them from influencing the repair-outcome predictions.

### 2.6. Handling variable output classes

To handle the varying Δlength classes reported across datasets, we define a global output range comprising 188 classes (−99 to +88 bp). We address the discrepancy between this global range and the narrower, dataset-specific output classes by constructing a binary mask, *M*_*d*_ ∈ {0, 1}^188^, for each dataset *d*. This mask designates the subset of Δlengths explicitly measured in a given experiment. During training, the loss is computed over the dataset-specific subset of classes; specifically, CROP’s predicted distribution is renormalized over the indices where *M*_*d*_ = 1 prior to evaluating the loss. This approach ensures the model is not penalized for unobserved classes, while enabling the learning of a cohesive representation across datasets.

### 2.7. Training CROP

To improve prediction robustness, we trained an ensemble of 9 independent models and averaged their predicted Δlength probability distributions during inference. The training procedure for each model was divided into two distinct phases. In the first phase, we trained all model parameters for 50 epochs with Kullback–Leibler divergence (KLD) loss, Adam optimizer (default parameters: *β*_1_ = 0.9, *β*_2_ = 0.999), a learning rate of 10^−3^, and a batch size of 512. In the second phase, to refine the dataset-specific embedding without altering the learned sequence logic, we froze all weights aside from the dataset embedding vectors and trained for an additional 10 epochs with the same hyper-parameter. All hyper-parameter were selected based on preliminary testing rather than extensive hyperparameter optimization, since preliminary FSR prediction performance on the validation set was already better than competing methods’. We trained the models on the Linux server described in Section 2.3. The training of all ensemble models on all datasets terminated in less than 12 hours.

To demonstrate that CROP did not overfit within those 60 epochs, we report the validation loss and validation FSR prediction performance during training, using a the same training data and hyper-parameters as in Section 2.7, on a validation set we held out from the training set (Supplementary Section S5).

### 2.8. Ablation analysis

To test the contribution of the MH module and the dataset-specific embedding on CROP’s performance, we evaluated the performance of CROP in 9-fold cross-validation on all data outside the test set. For runtime purposes, for each partition, we did not train an ensemble, but rather trained a single model. We quantified the impact of these components by evaluating the change in Pearson correlation under two configurations: (i) the exclusion of the MH module, and (ii) the removal of the dataset-specific embedding.

### 2.9. Interpretability analysis

#### 2.9.1. In-silico mutagenesis

To quantify the contribution of individual nucleotides to a specific Δlength class *y*, we applied *in-silico* mutagenesis (ISM). For a given target sequence *X* of length *L*, we systematically mutate the nucleotide at position *i* to every alternative base *b*^*′*^ ∈ {*A, C, G, T*} and compute the change in the predicted probability for the class *y*. The mutation effect *S*(*X*)_*i,b ′*_ is defined as:

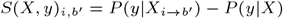

To map the nucleotide base *b*^*′*^ to an index *j*, we use a mapping function *f* : {*A, C, G, T*} *→* {1, 2, 3, 4}. Thus, for each position *i* and nucleotide index *j* we create an *L* × 4 mutation map:

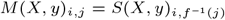

The mutation map thus captures which specific mutations increase or decrease the likelihood of class *y* relative to the wild-type target sequence. To plot the sequence logos, we mean normalize *S*(*X, y*)_*i,b′*_ per position *i*.

#### 2.9.2. Pairwise interaction scores

To capture pairwise interactions, with respect to a specific Δlength class *y*, where the importance of position *t* is dependent on the identity of position *s*, we developed a technique we call *pairwise interaction scores* (PIS), which is inspired by DFIM [Greenside et al., 2018]. PIS measures the variation of the ISM profile under perturbations.

Let **M**(*X, y*) be an *L* × 4 mutation map for target sequence *X* and class *y*. To measure the influence of a “source” position *s* on a “target” position *t*, we mutate *s* to base 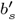 and recalculate the mutation map, denoted as 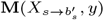. The dependency score 𝒟 (*X, y*)_*s,t*_ represents the aggregate change at *t* caused by the mutation at *s*:

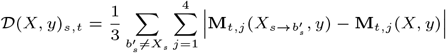

The resulting *L* × *L* map *D*(*X, y*) reveals pairwise interactions, where high values at *D*(*X, y*)_*s,t*_ indicate that position *t* is dependent on position *s*, with respect to class *y*.

#### 2.9.3. In-silico microhomology perturbations

To systematically quantify how MH affects FSR, independent of confounding sequence contexts, we utilized CROP as an *insilico* experimental oracle, using a technique we call *in-silico* MH perturbations (IMP), which was inspired by GIA [Koo et al., 2021]. We generated random DNA backgrounds *X*_*syn*_ embedded with repetitive sequences of varying lengths (*L*_MH_) placed symmetrically at varying gaps (*d*_gap_) upstream and downstream of the canonical cut site. We enforced the presence of the PAM sequence (NGG) at the expected position; if an embedded sequence downstream of the canonical cut site overlapped with the PAM, the PAM was preserved and correctly mirrored to the upstream embedded sequence. We implemented a flanking-nucleotide constraint that prevented the accidental formation of repetitive sequences longer than *L*_MH_.

We define the expected MMEJ deletion as Δ_MMEJ_(*d*_gap_, *L*_MH_) = −(2 · *d*_gap_ + *L*_MH_), representing the excision of the symmetric gap length (2 · *d*_gap_) along with the length of the repetitive sequence (*L*_MH_). We approximated the probability of the expected MMEJ deletion:

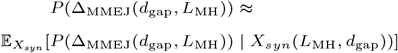

For all approximations, we used 200 generated sequences as a tradeoff between accuracy and runtime.

## 3. Results

### 3.1. CROP outperforms state-of-the-art methods in frameshift prediction

We trained CROP on all datasets but the SPROUT dataset, which we left as an independent held-out test set, following the evaluation protocol of CROTON and Apindel. We gauged prediction performance by Pearson correlation between measured and predicted FSR over each dataset separately (Figure 3). For SPROUT^CA^, we evaluated CROP by using the FORECasT K562 dataset embedding and mask, as it was the only dataset which multiple models (FORECasT, CROTON, and Apindel) trained on.

**Fig. 3.**
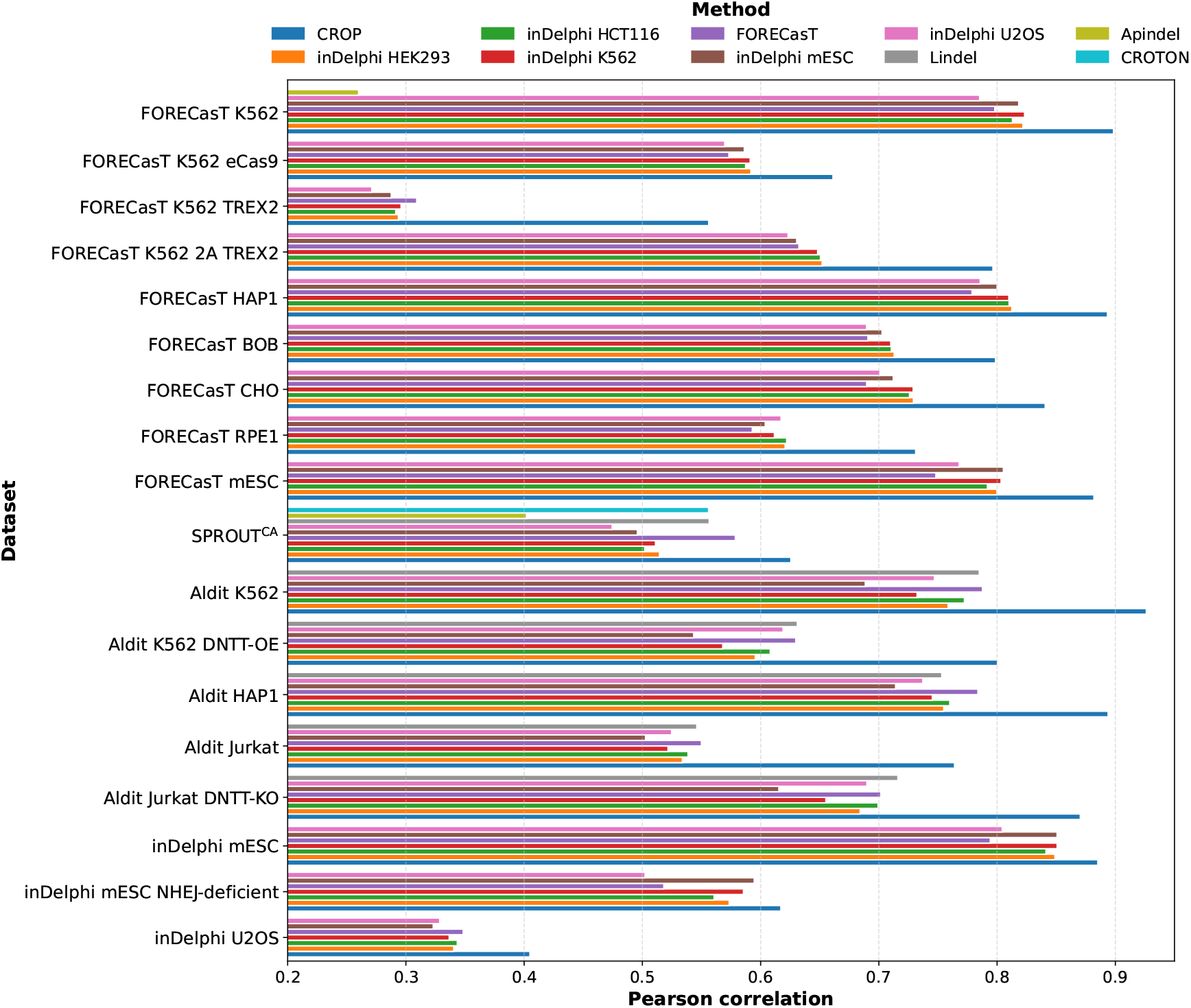
Prediction performance of CROP compared to publicly available methods for FSR prediction. We gauged prediction performance by Pearson correlation of measured and predicted FSR. For SPROUT^CA^ (which CROP was not trained on), we generated CROP’s predictions using the FORECasT K562 dataset embedding and mask. We ordered methods by the number of datasets they can be evaluated on, and then by their achieved average correlation over all the datasets they were evaluated on.

CROP achieved superior FSR prediction performance compared to all other publicly available methods over all datasets (Wilcoxon rank-sum test *p <* 0.006 for methods where predictions were available for all 18 datasets, and *p <* 0.03 for Lindel which was available for only 6 datasets due to minimum sequence length requirements). Notably, CROP generalized better on SPROUT^CA^, a dataset which none of the methods trained on, achieving a Pearson correlation of 0.626 compared to the second best model, FORECasT, achieving 0.579. CROP improved prediction performance over unique datasets, i.e., Aldit Jurkat, and Aldit K562 DNTT-OE, where an abundance of DNTT resulted in increased insertion events [Zhang et al., 2023], and the FORECasT K562 TREX2 dataset, by large margin: from the second best method achieving [0.550, 0.631, 0.309] to [0.764, 0.800, 0.556] by CROP, respectively.

To compare CROP to CROTON and Apindel (as we couldn’t run them) on the FORECasT K562 dataset on which both were trained on, we retrieved the AUROC values reported in their studies. The AUROC values were calculated by labeling FSR above the median as positive. CROP achieved an AUROC of 0.927 compared to CROTON’s 0.800 and Apindel’s 0.690, thus outperforming their reported FSR prediction performance on on the FORECasT K562 dataset.

### 3.2. Datasets concordance and context transferability

We sought to evaluate the concordance of CROP’s dataset-specific embeddings across diverse datasets. To this end, we performed a systematic cross-evaluation where the FSRs for each test set were predicted using the dataset-specific embedding and mask associated with every dataset (Figure 4A).

**Fig. 4.**
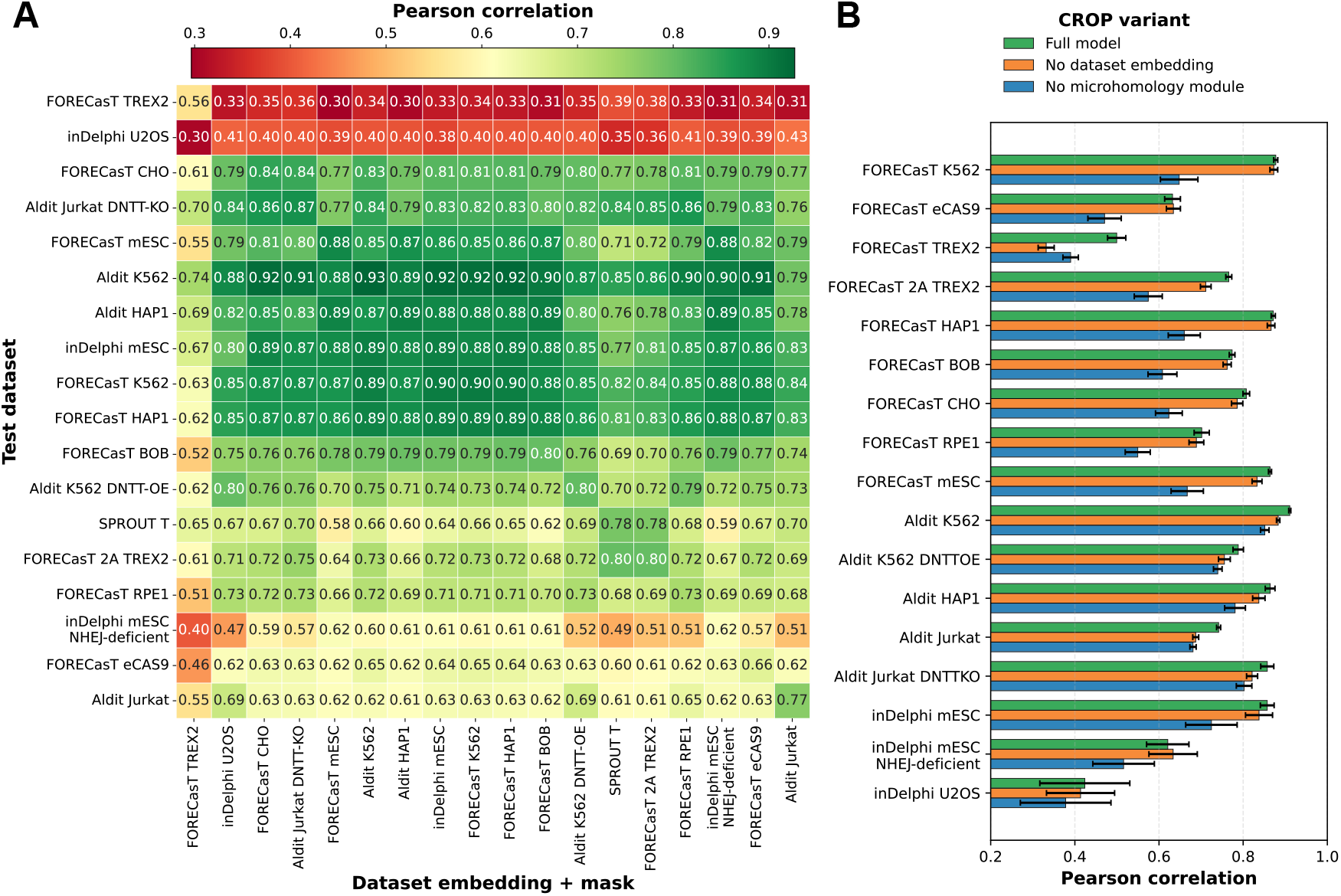
Dataset concordance and ablation analysis. (A) Concordance between experimental datasets. We report the Pearson correlation between measured and predicted FSR, achieved by CROP trained on all datasets excluding SPROUT^CA^. Columns indicate the dataset embedding and mask used in inference, rows indicate the test dataset used. Rows and columns were reordered using hierarchical clustering (UPGMA) to group test datasets that show similar performance across different embeddings and masks. (B) Ablation analysis. The effects of removing each of the microhomology module and dataset embedding, separately, from CROP. We performed 9-fold cross validation on the original training set (without using an ensemble, for time purposes). We report the mean Pearson correlation between measured and predicted FSRs on the validation folds.

CROP was trained and tested on all datasets excluding SPROUT^CA^ with train-test partitioning as described in Section 2.2.

In most cases, cross-embedding resulted in a Pearson correlation *≥* 0.73, suggesting a high degree of shared repair principles. However, certain datasets (i.e., FORECasT TREX2, inDelphi U2OS, and inDelphi mESC NHEJ-deficient) exhibited distinct distributional shifts, leading to lower performance when using other datasets’ embedding. Interestingly, the dataset tested proved had a greater effect on prediction performance than the dataset-specific embedding or mask. For instance, on the inDelphi U2OS dataset the Pearson correlations ranged from 0.3 to 0.43, while for the Aldit K562 dataset, the correlations ranged from 0.74 to 0.93.

The Aldit K562 embedding may serve as a robust default, achieving the highest mean Pearson correlation (0.736 *±* 0.167). While numerically superior, this performance was not significantly different compared to 16 other tested embeddings (pairwise Wilcoxon rank-sum tests, Benjamini-Hochberg adjusted p *>* 0.05). A significant reduction in performance was only observed when compared to the FORECasT TREX2 embedding, which achieved a mean of 0.578 *±* 0.109 (adjusted p = 0.024).

### 3.3. Ablation analysis of the microhomology module and dataset embedding in CROP

We performed an ablation analysis on the same training set as in Section 3.1 with partitioning to training and validation sets as described in Section 2.8. The analysis demonstrated that both the MH module and dataset embedding contribute to CROP’s improved performance. CROP achieved a mean Pearson correlation of 0.756*±*0.140, representing a performance ceiling for FSR prediction (Figure 4B).

Removing the MH module resulted in a performance degradation across all 17 datasets, with the mean Pearson correlation dropping to 0.627 *±* 0.136. Similarly, excluding dataset embeddings reduced average correlation to 0.727 *±* 0.156 (adjusted *p <* 0.001 in both comparisons by pairwise Wilcoxon signed-rank test with Benjamini-Hochberg adjustment). Interestingly, while the dataset embedding improved performance in 15 of the 17 datasets, we observed exceptions in the FORECasT eCAS9 and InDelphi mESC NHEJ-deficient datasets. In those two datasets, removing dataset embedding resulted in a non-statistically significant increase in Pearson correlation from 0.632 *±* 0.019 and 0.621 *±* 0.050 to 0.634*±*0.017 and 0.634*±*0.058 (*p >* 0.05), respectively.

### 3.4. CROP learns microhomology principles

To interrogate CROP for the repair principles it learned, we visualized sequence preferences and nucleotide interactions using ISM (Section 2.9.1) and PIS (Section 2.9.2). For Δlength ∈ {−5, −7, −10, −15, −20}, we analyzed the three target sequences with the highest experimentally measured probability for that event in the FORECasT K562 dataset (Figure 5 and Supplementary Section S6). We chose these target sequences, as we expected them to exhibit MH-associated patterns.

**Fig. 5.**
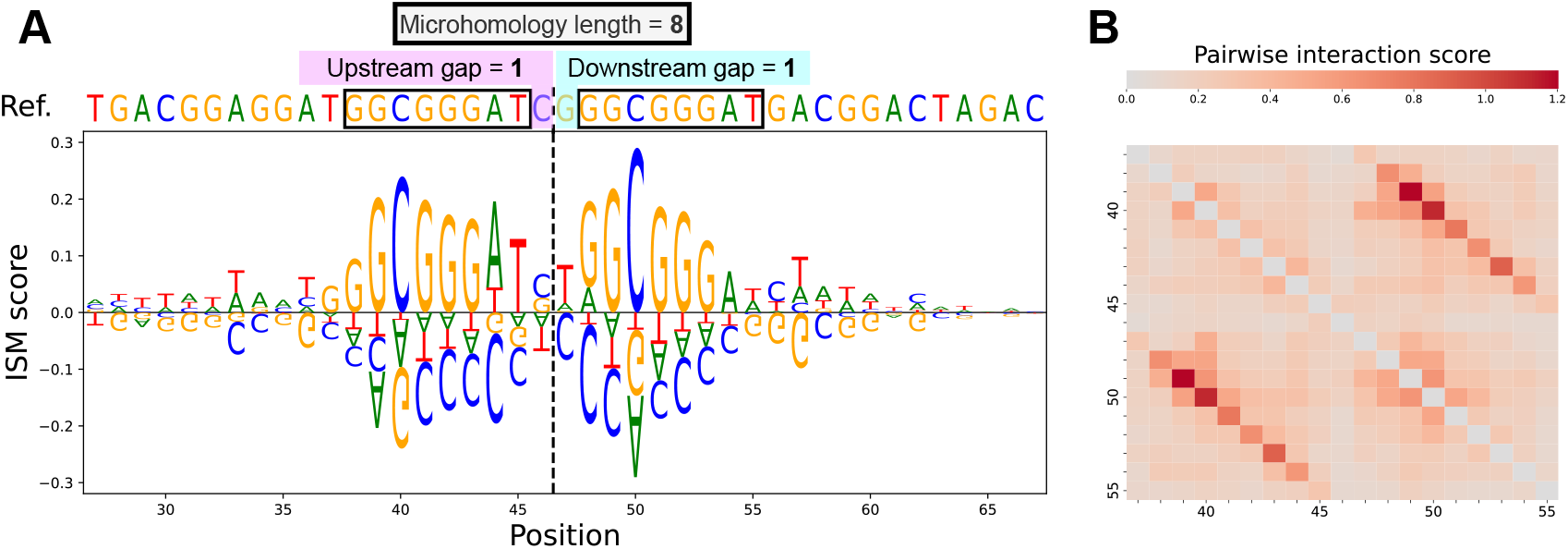
Interpretability analysis of a representative target sequence for Δ = −10 in the FORECasT K562 dataset, centered and cropped around the cut site. (A) In-silico mutagenesis profile, where the height of each nucleotide represents its mean-shifted effect on CROP predicting the Δ = −10 class. (B) Pairwise interaction scores to indicate pairwise position dependence with respect to CROP predicting the Δ = −10 class.

The resulting ISM profiles reveal that CROP assigns high importance to the repetitive sequences flanking the canonical cut site. Furthermore, PIS indicate that the nucleotides within these repetitive sequences interact with one another as evident by the stretches parallel to the diagonal in Figure 5B and Supplementary Section S6. These results demonstrate that CROP successfully learns MH principles directly from target sequences without requiring explicit feature engineering.

We further investigated CROP’s learned repair principles using IMP (Section 2.9.3) over MH length *L*_MH_ ∈ [1, 20] and symmetric gap distance *d*_gap_ ∈ [0, 10] using the FORECasT K562 dataset embedding and mask, to explore their effect on predicting the probability of the expected MMEJ deletion Δ_MMEJ_(*d*_gap_, *L*_MH_) = −(2 *d*_gap_ + *L*_MH_) (Figure 6). Our investigation revealed that for a fixed expected MMEJ deletion, CROP predicts a higher probability when utilizing longer repetitive sequences paired with shorter gaps. Furthermore, our investigation revealed that, generally, as MH length increases, the probability of the expected MMEJ deletion is increased, hinting at a possible preference for the the MMEJ pathway. Finally, while the probability of the expected MMEJ deletion generally increases with *L*_MH_, we observe an eventual decrease in the probability for symmetric gap distances *≥* 4; we hypothesize this trend reflects data limitations, as the training data may under-report or miss long genomic deletions.

**Fig. 6.**
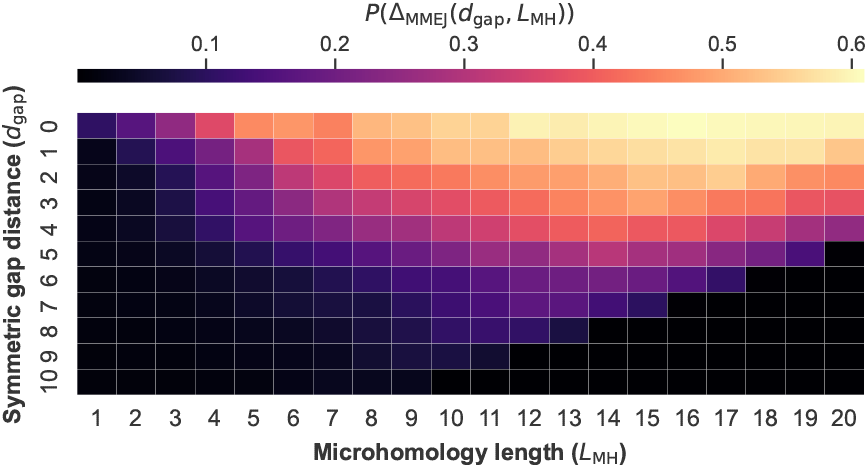
Probability of the expected MMEJ deletion Δ_MMEJ_(*d*_gap_, *L*_MH_) estimated using *in-silico* microhomology perturbations on the FORECasT K562 dataset embedding and mask, over microhomology length *L*_MH_ ∈ [1, 20] and symmetric gap distance *d*_gap_ ∈ [0, 10].

## 4. Discussion

In this study, we conducted the most thorough FSR prediction tests to date, spanning 18 datasets and 10 methods to assess whether state-of-the-art methods are able to predict FSR accurately over current datasets. We curated datasets from four studies and standardized them to Δlength format, enabling future studies to access repair-outcome datasets without prior knowledge of each study’s preprocessing methodology.

In addition, we developed CROP using cellular-context embedding and no MH features to train over these datasets. CROP outperformed all state-of-the art methods in FSR prediction over all datasets. Our ablation analysis confirmed that the MH module and dataset embedding contribute to CROP’s performance.

Our interpretability analysis using ISM and PIS confirmed that CROP learned MH-mediated repair mechanisms directly from target sequences. Furthermore, our IMP analysis showcased how CROP’s predictions are affected by MH length and symmetric gap distance.

There are several limitations to this study. First, we rely on preprocessing performed by the original studies as our starting point, which do not follow the exact same protocol. This may be a cause for discrepancies between datasets, and lead to suboptimal models. Second, for a specific dataset, CROP is limited to predict only on Δlength observed in the original dataset, missing unobserved Δlength ranges. Third, we did not conduct an extensive hyperparameter and architectural search, which may improve FSR prediction even further.

We have several plans to improve CROP in the future. First, we plan to re-process all datasets from raw reads whenever available to adhere to a unified preprocessing protocol, thus removing some datasets discrepancies. Second, we plan to experiment with various MH modules, e.g., pre-trained on sequence similarity tasks. Third, we will interrogate CROP further, in particular, in regards to staggered cuts and the representation of each cellular context. Finally, we plan to make CROP accessible via a user-friendly web server.

## 5. Conclusion

In conclusion, we introduced CROP, a new state-of-the-art method to predict FSR across various cellular contexts. Our study suggests that CROP learned the MH mechanism, which may have contributed to its generalization. CROP can be used in gene knockout applications to find target sequences with high FSR following a DSB, thus leading to improvement in gene-editing applications.

## Acknowledgments

I.T. acknowledges the Bar-Ilan University Presidential Fellowship, the Planning and Budgeting Committee Data Science PhD Fellowship, and the cloud computing credit by the Israel Data Science and AI Initiative.

## Data availability

The processed Δlength datasets, alongside test splits used in evaluating CROP, are archived on Figshare at https://doi.org/10.6084/m9.figshare.30998056.

