## Supplemental Information for "CROP: A feature-independent context-aware method for CRISPR-Cas9 frameshift prediction"

### S1. X-CRISP's FORECasT preprocessing

We evaluated the compatibility between the original FORECasT datasets and those preprocessed by X-CRISP. We report the Pearson correlation for each observed Δlength (Figure S1). The FSR Pearson correlation for HAP1, TREX2 and mESC was 0.775 0.395 and 0.786, respectively. One major reason for this disparity is that datasets preprocessed X-CRISP do not contain mixed events.

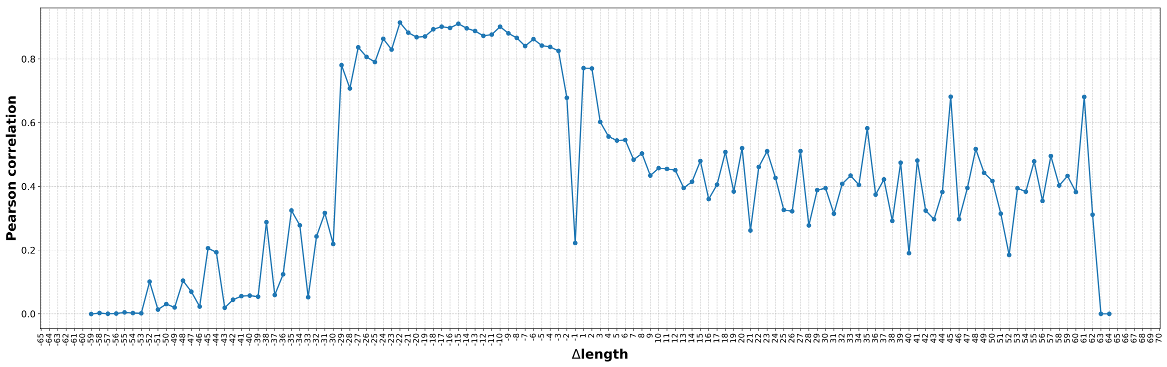


FORECasT HAP1

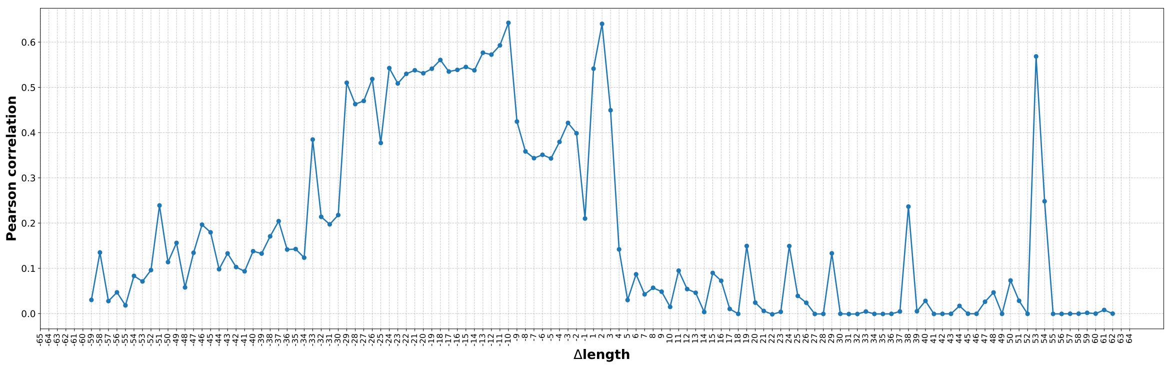


FORECasT TREX2

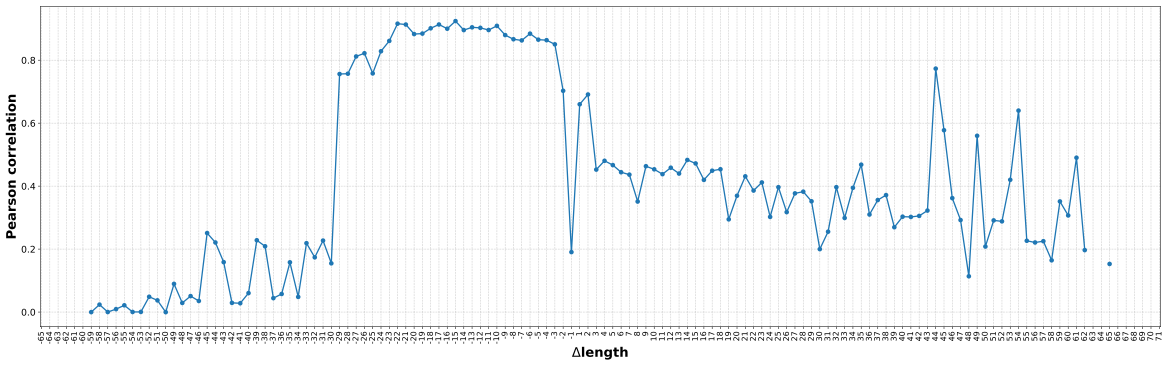


FORECasT mESC

**Fig. S1.** Pearson correlation of varying $\Delta$length values, between X-CRISP's preprocessing and FORECasT's preprocessing of the same FORECasT datasets.

FORECasT datasets preprocessed by X-CRISP exhibit a different $\Delta$length distribution than the FORECasT's and contain a spike for $\Delta\text{length} = -1$ (Figure S2).

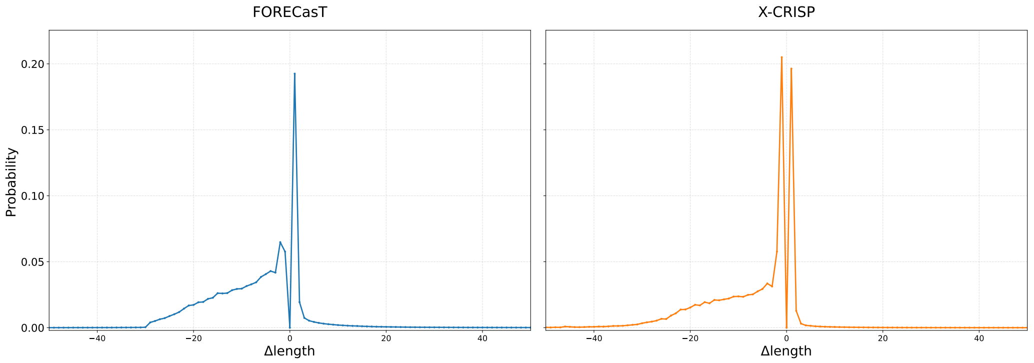


FORECasT HAP1

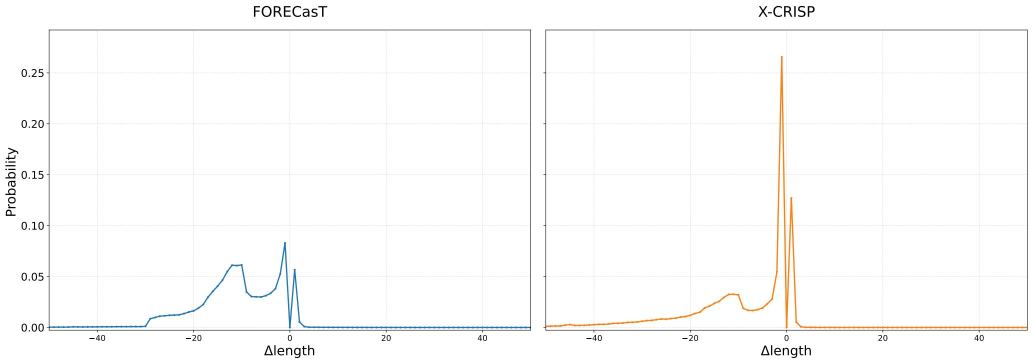


FORECasT TREX2

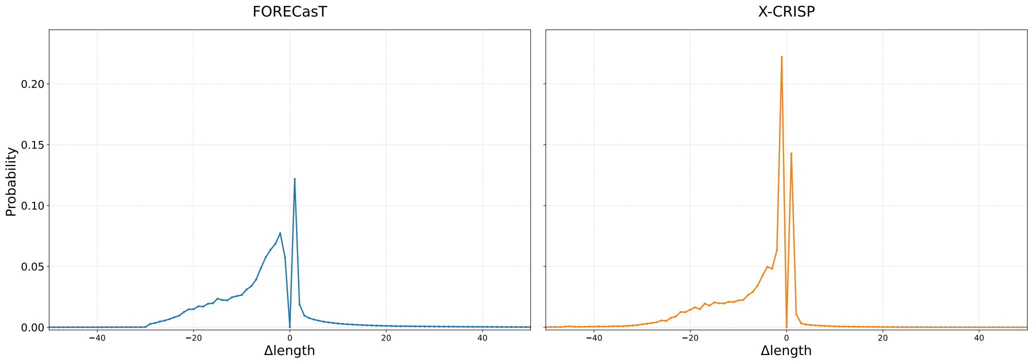


FORECasT mESC

**Fig. S2.** Comparison of $\Delta$length probabilities calculated after summing reads over all target sequences, between the original FORECasT study's preprocessing and X-CRISP preprocessing across FORECasT HAP1, FORECasT TREX2, and FORECasT mESC datasets, within the range of $\Delta\text{length} \in [-50, 50]$.

### S2. Curating datasets

#### S2.1. FORECasT

We utilized publicly available repair outcomes from Figshare [Figshare] and Supplementary Data 1 from the study [Paper], which provided the target sequences. To generate each dataset, we processed the CIGAR strings (e.g., `D19_L-21I1C2R1`) to extract Δlengths, omitting the provided mutated sequences. This approach was taken to replicate CROTON's and Apindel's preprocessing.

We selected replicates that maintained 800x coverage and were harvested 7 days post-infection. While both repetitions were included for most of the dataset, the FORECasT RPE1 sample was limited to a single repetition which only had 500x coverage.

#### S2.2. SPROUT

We used SPROUT's files from Figshare [Figshare], making sure to remove mixed events and insertions longer than 20 bp for $\text{SPROUT}^{\text{CA}}$. While most papers report 1,603 target sequences, this is partially correct, as out of the 1,603 target sequences we were able to retrieve, only 1,598 of them have any reported indels, effectively meaning that the observed on-target efficiency was 0.

#### S2.3. Aldit

We sourced our Aldit datasets - `GSE181774_DSB_Repair_Map.xlsx` from Gene Expression Omnibus [GEO]. For the K562 DNTT-OE and Jurkat DNTT-KO datasets where a target sequence wasn't present in the data, we tried to match the column "sgRNA_name" to any of the other Aldit datasets in order to retrieve the target sequence. In cases where we couldn't find a match, we discarded the sample. Afterwards we discarded all columns representing an unknown amount of insertion or deletions, such as "I-3+" or "D30+", before converting to a Δlength format.

#### S2.4. inDelphi and X-CRISP

All inDelphi datasets were taken from X-CRISP's preprocessing, archived on Figshare [Figshare]. We used this same source for XCRISP's preprocessing of FORECasT.

#### S2.5. Lindel

We considered using the dataset from Figshare [Figshare]. Opening `Lindel_training.txt`, we saw that it reports the 20nt-long guide RNA, without the PAM. We considered this dataset insufficient for modeling MH-mediated deletions, and did not use it in our study.

### S3. Methods which we could not benchmark

#### S3.1. CROTON

Following the procedures outlined in CROTON's repository [GitHub page], we attempted to initialize the `conda` environment using the provided `croton.yml` file. However, environment creation failed due to a `PackagesNotFoundError`, indicating that several dependencies were unavailable in the specified conda channels.

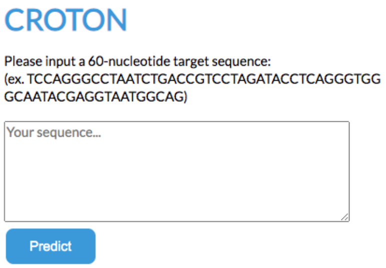


**Fig. S3.** CROTON's interface, taken from its GitHub page.

Furthermore, the publicly available implementation of CROTON is structured as a self-hosted `Django` web server designed for single target sequence predictions (Figure S3), which is infeasible for our benchmarks. Upon contacting the authors, they confirmed that no standalone batch-processing script or command-line interface was available to facilitate large-scale evaluation outside of the local web server framework.

#### S3.2. Apindel

We attempted to evaluate Apindel using the documentation provided in its GitHub repository [GitHub page]. The authors do not provide model weights, rather suggesting that users reproduce results by integrating the FORECasT, SPROUT, and Lindel datasets using the code provided in the `model_creation` directory.

However, we found that the repository lacks documentation detailing the required preprocessing steps to transform these raw datasets into the model's expected input format, with no library versions specified throughout the repository. Consequently, we were unable to reconstruct Apindel for out benchmarks. We have tried reaching out to the authors of Apindel, and as of now have received no reply.

#### S3.3. Aldit

While CRISPR-Aldit was originally hosted on GitHub [GitHub page], the repository now directs users to a web-based interface [Web page]. According to the repository notice the source code and pre-trained models were migrated to this external platform.

However, we were unable to use Aldit for benchmarks, as the web server frequently exhibited downtime and lacks a batch-processing feature. Despite reaching out to the authors to request access to the standalone models, we received no response as of the time of writing. Furthermore, a review of the repository's version history did not provide sufficient information to obtain and run the trained models.

#### S3.4. X-CRISP

We attempted to evaluate X-CRISP using the implementation provided in its GitHub repository [GitHub page]. Initial environment configuration using the provided `requirements.txt` failed, as the inclusion of `awscli` and `botocore` triggered an infinite recursion loop during the `pip` dependency resolution process. As these packages are primarily used for AWS service management and are not intrinsic to the model's architecture, we removed them to proceed with the setup.

However, executing the prediction script (`python predict.py`) resulted in a critical `ValueError` related to `numpy.dtype` size mismatch. This error indicates a binary incompatibility between the compiled C headers and the Python environment, a common issue when library versions are not strictly pinned. Because the repository's documentation and dependency files lacked specific version requirements for `NumPy` and other core libraries, we were unable to resolve the versioning conflicts, rendering the tool non-functional for our benchmarks. We contacted the authors of X-CRISP in order to obtain the correct versions required, but as of yet we have received no reply.

### S4. CROP's architecture

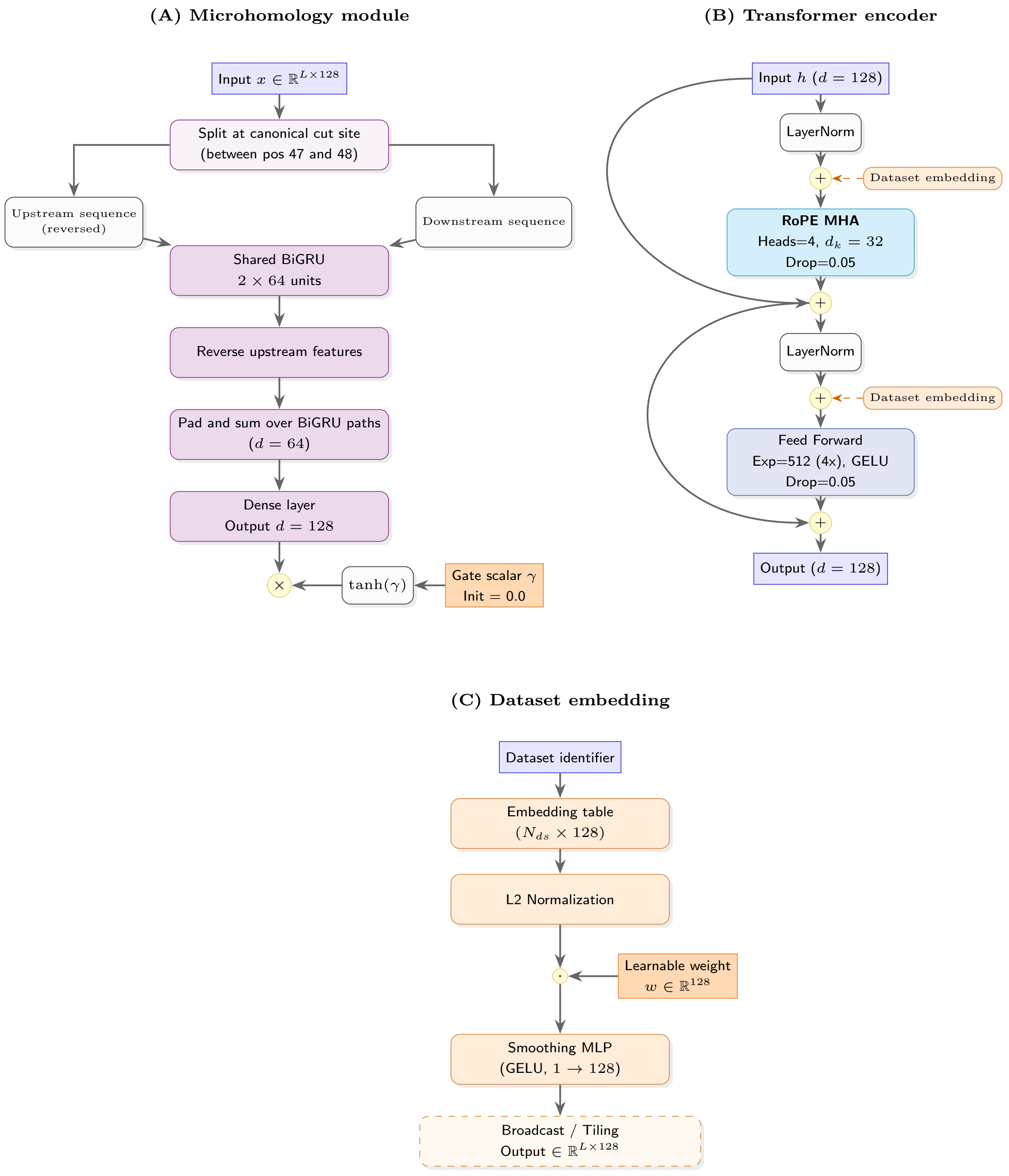


**Fig. S4.** A) Microhomology module. B) Transformer encoder block. C) Dataset embedding.

### S5. CROP's epoch count justification

To evaluate whether CROP overfits within the 60-epoch training window, we performed a performance analysis across the 17 experimental datasets used in the ablation study (Section 2.8). We gauged performance by Pearson correlation between measured and predicted FSR on the validation set, and by validation loss. We report the mean Pearson and mean validation loss, each calculated as the average performance of all individual models at each epoch.

Towards the end of training (epochs 40-60), the mean Pearson correlation generally trended upward or reached a plateau (Figure S5 and Figure S5). Overfitting was observed exclusively in the inDelphi U2OS dataset, beginning after approximately five epochs (Figure S7). However, given that the Pearson correlation continued to improve across all other datasets beyond this point, and that the overall validation loss has not increased (Figure S8) we opted to prioritize global improvements over the specific optimization of the inDelphi U2OS dataset.

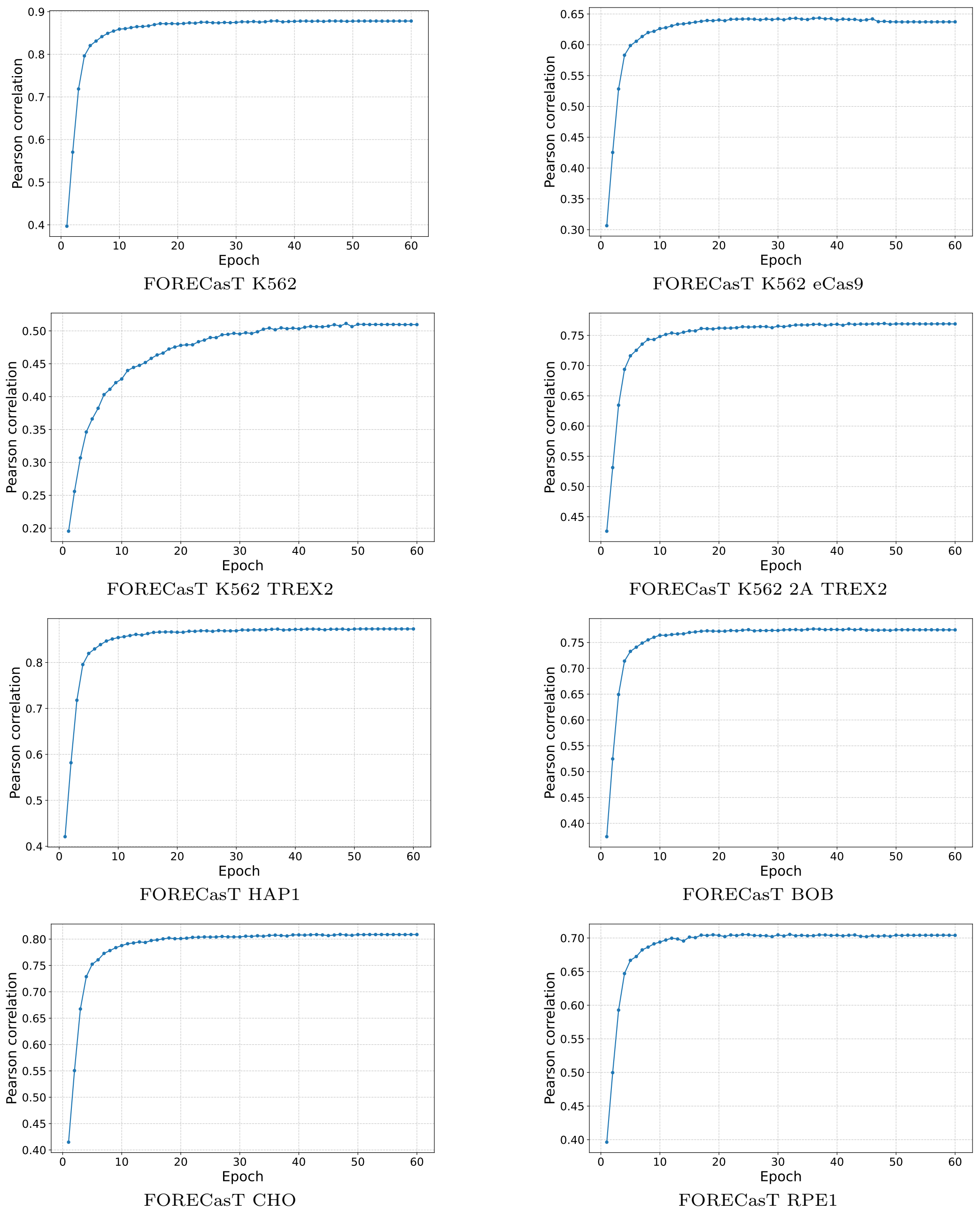


**Fig. S5.** Pearson correlation between measured and predicted FSR on the validation set. Values reported are averaged over 9 folds.

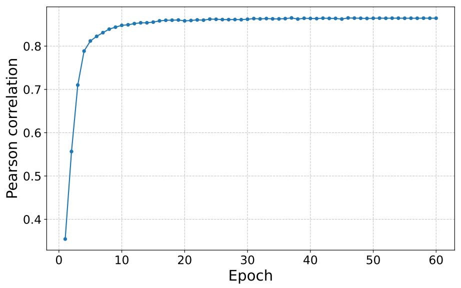


FORECasT mESC

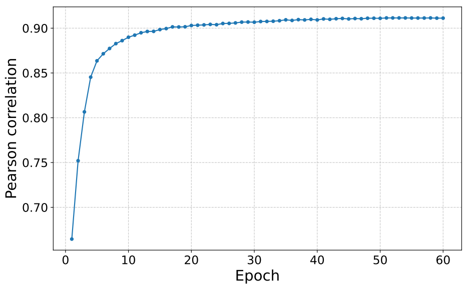


Aldit K562

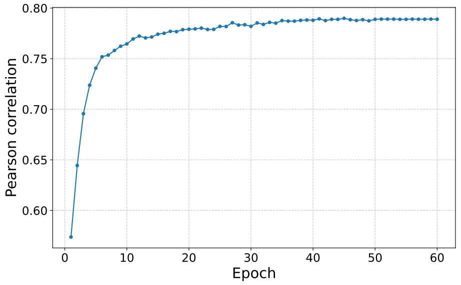


Aldit K562 DNTT-OE

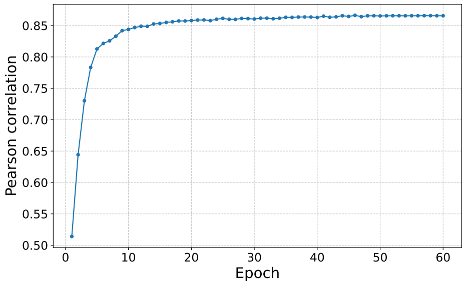


Aldit HAP1

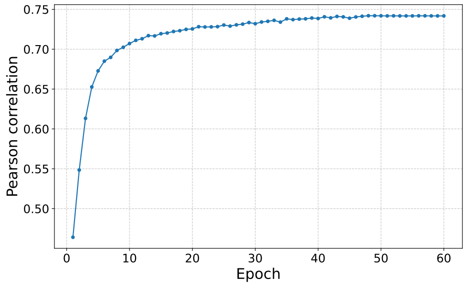


Aldit Jurkat

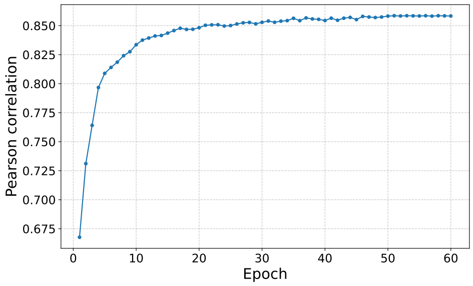


Aldit Jurkat DNTT-KO

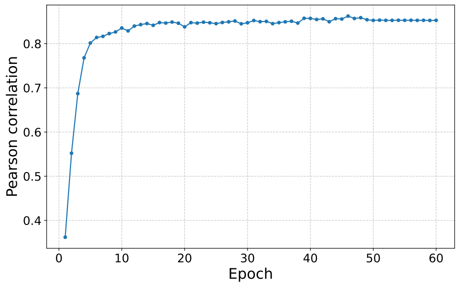


InDelphi mESC

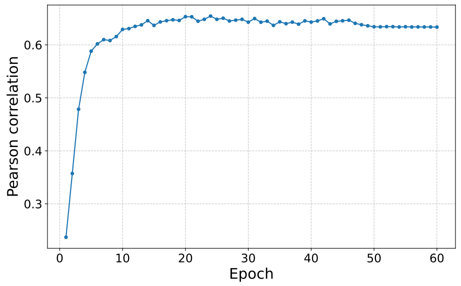


InDelphi mESC NHEJ-def.

**Fig. S6.** Pearson correlation between measured and predicted FSR on the validation set. Values reported are averaged over 9 folds. (cont.).

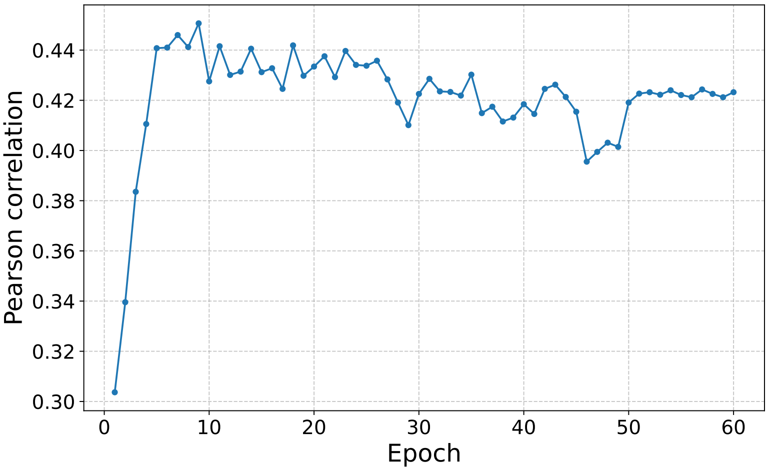


**Fig. S7.** Pearson correlation between measured and predicted FSR on the validation set, specifically, samples from the inDelphi U2OS dataset. Values reported are averaged over 9 folds.

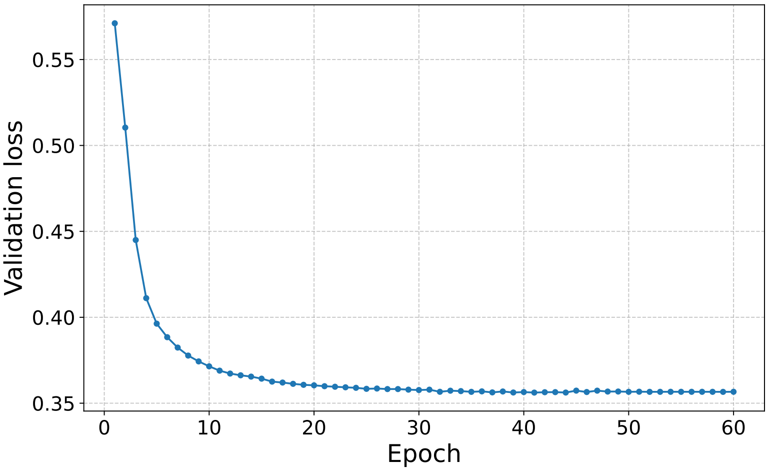


**Fig. S8.** Mean validation loss across training epochs, averaged over 9 folds.

### S6. In-silico mutagenesis profiles and pairwise interaction scores examples

This section presents the interpretability analysis for representative target sequences selected from the FORECasT K562 dataset. For $\Delta\text{length} \in \{-5, -7, -10, -15, -20\}$, we analyzed the three target sequences with the highest experimentally measured probability for that event in the FORECasT K562 dataset. The results are visualized using ISM (Section 2.9.1) and PIS (Section 2.9.2).

#### $\Delta$length $= -5$

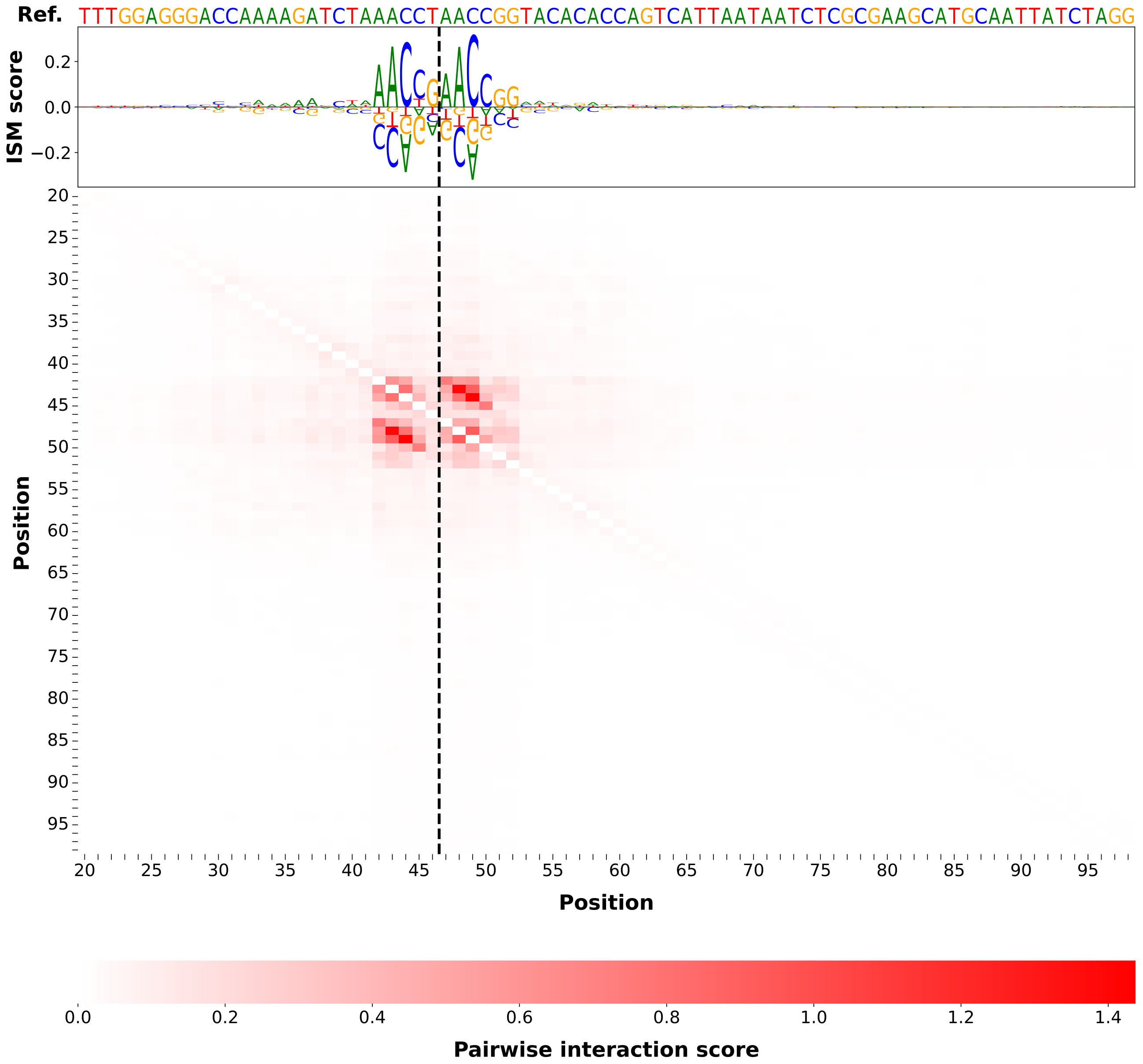

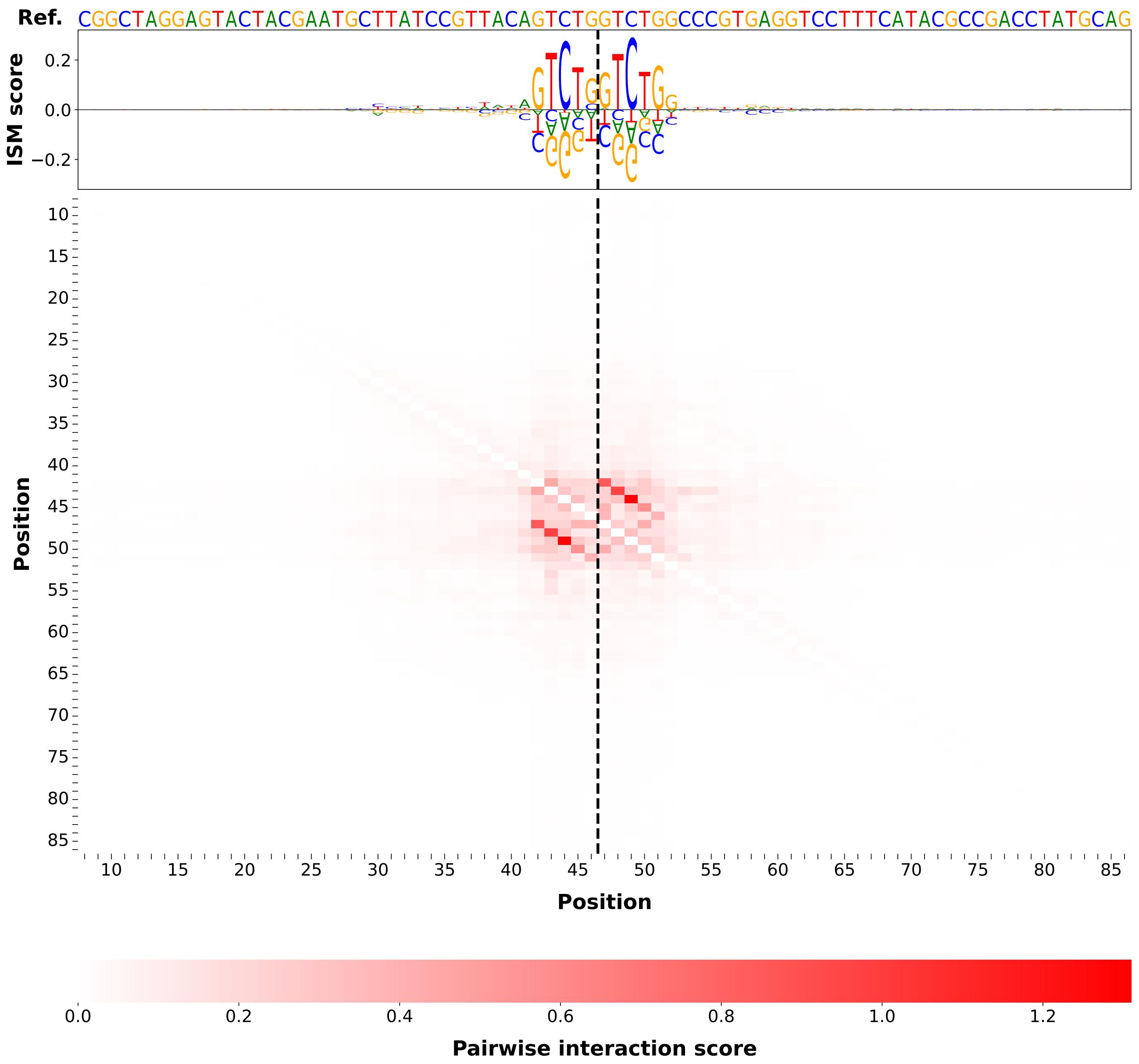
Ref. CGGCTAGGAGTACTACGAATGCTTATCCGTTACAGTCTGGTCTGGCCCGTGAGGTCCTTTCATACGCCGACCTATGCAG
ISM score
0.2
0.0
−0.2
Position
10
15
20
25
30
35
40
45
50
55
60
65
70
75
80
85
Position
0.0
0.2
0.4
0.6
0.8
1.0
1.2
Pairwise interaction score

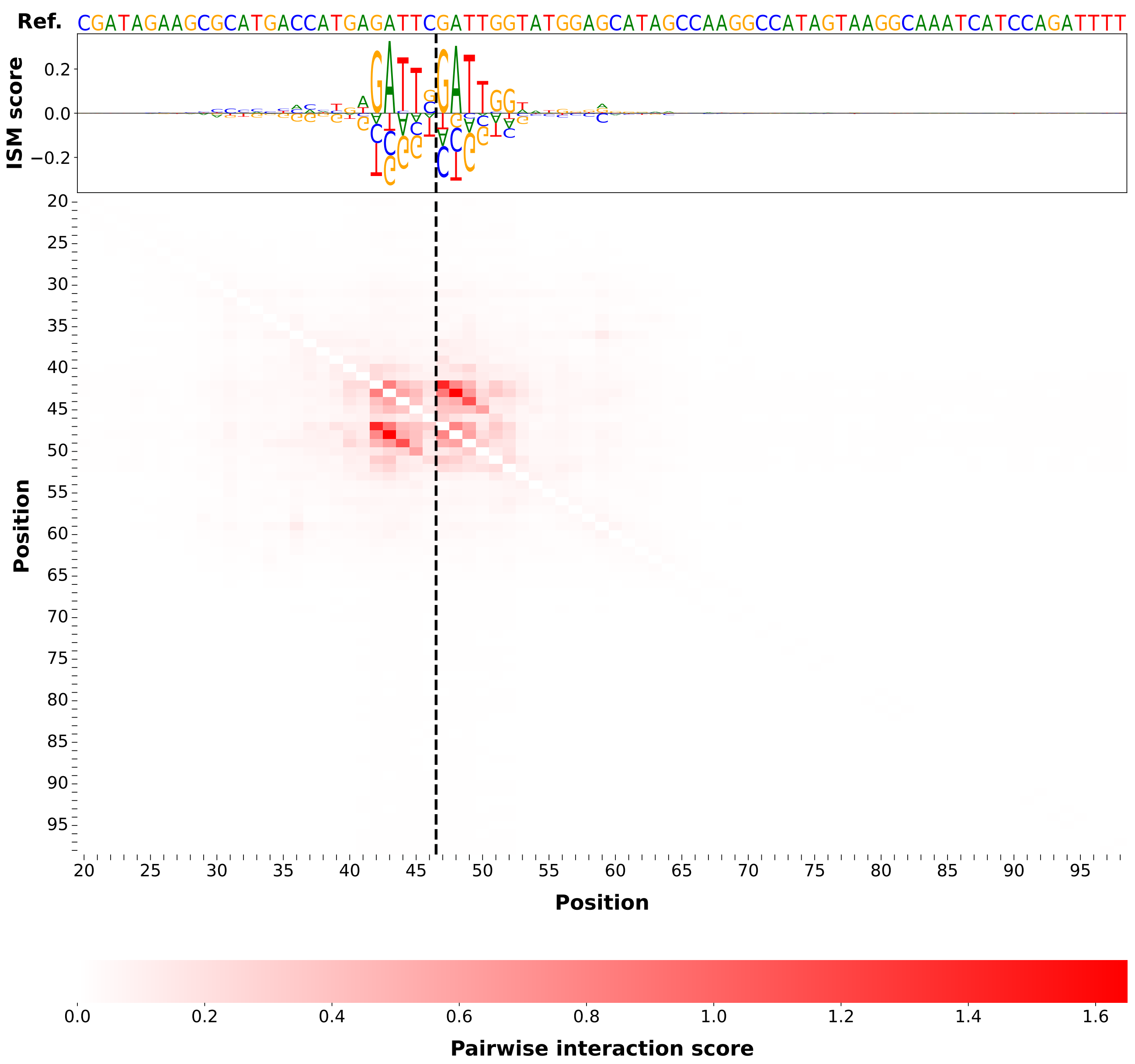
Ref. CGATAGAAGCGCATGACCATGAGATTCGATTGGTATGGAGCATAGCCAAGGCCATAGTAAGGCAAATCATCCAGATTTT
ISM score
Position
Position
Pairwise interaction score

$\Delta \mathrm{length} = -7$

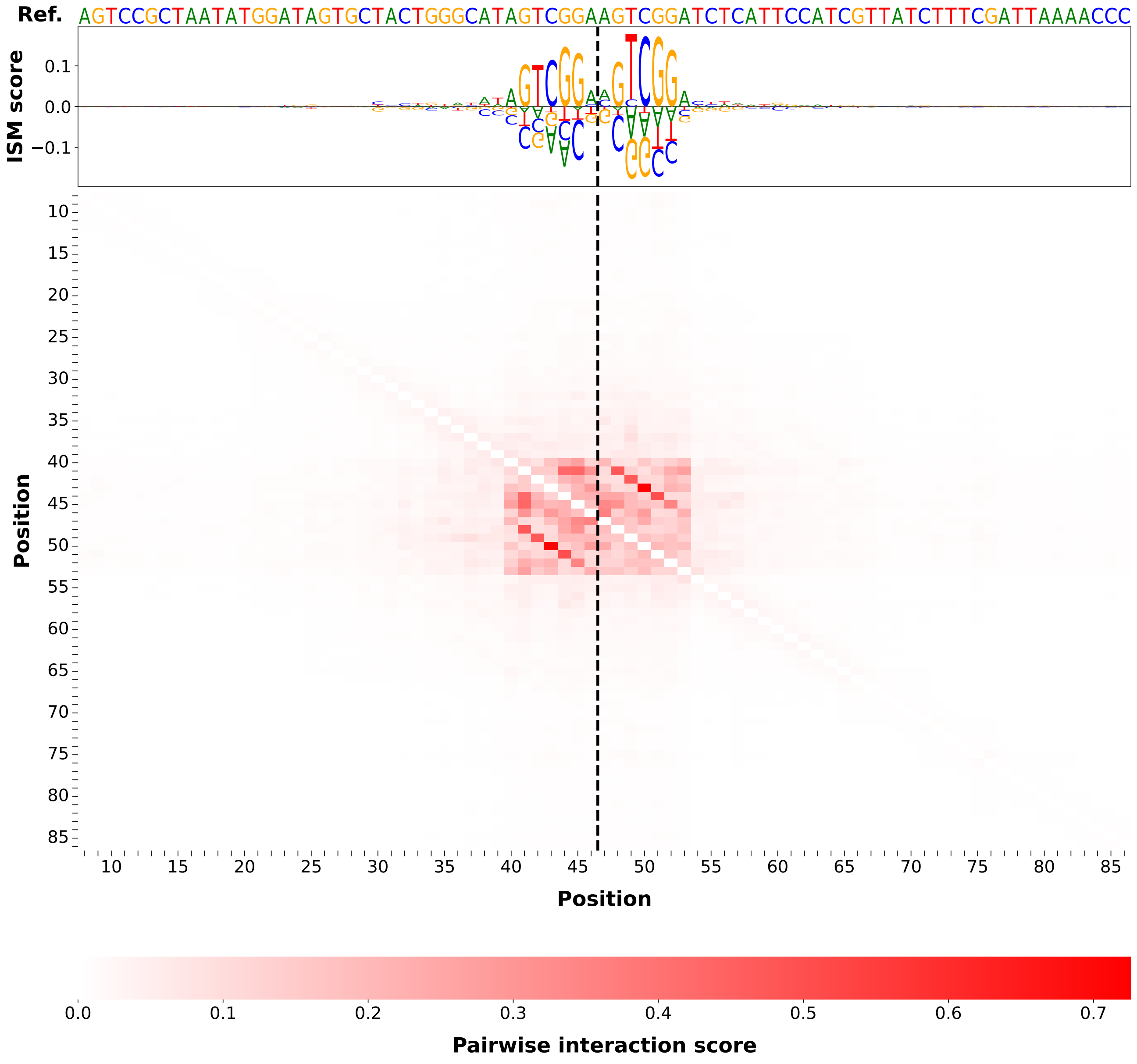

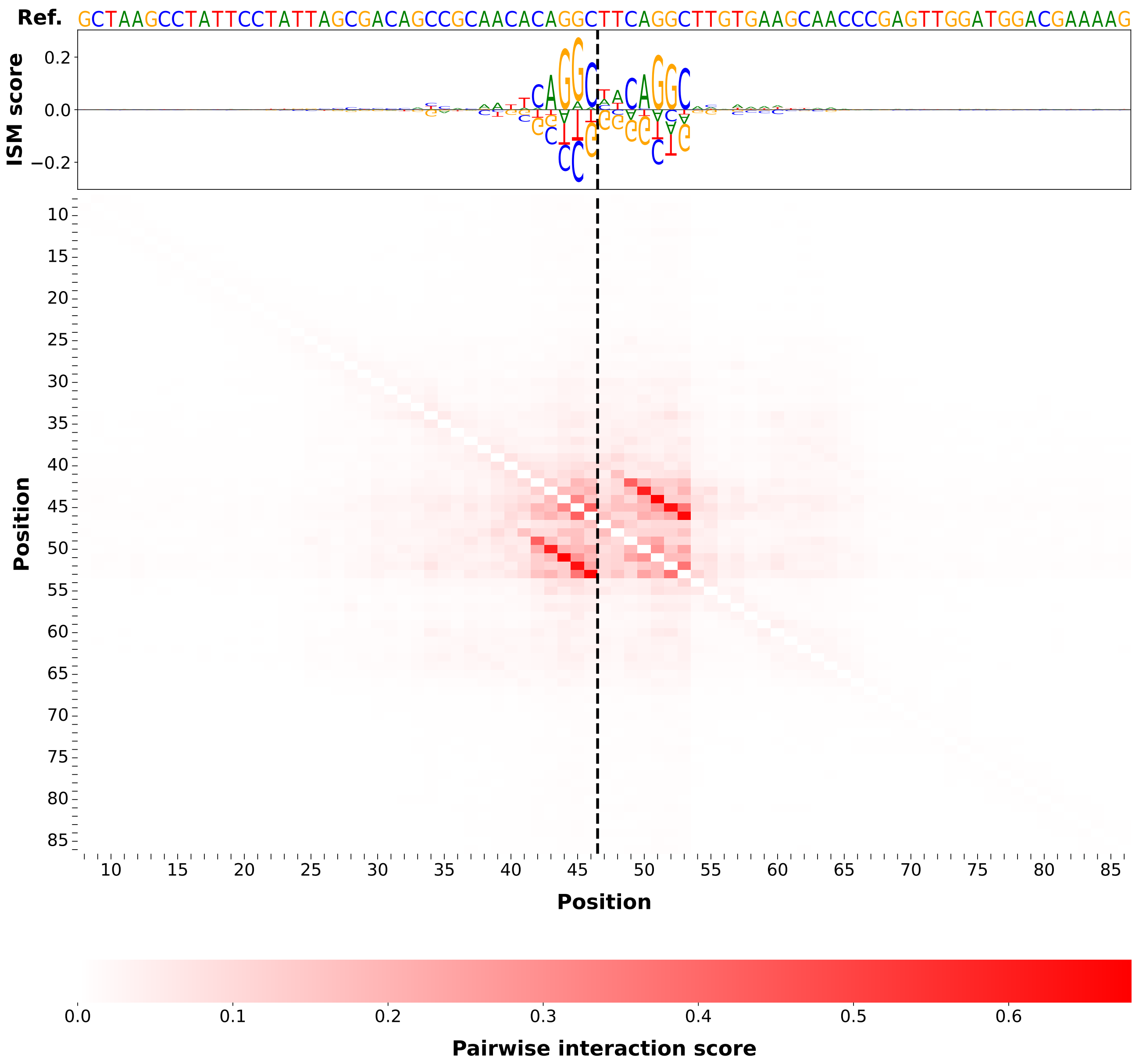
Ref. GCTAAGCCTATTCCTATTAGCGACAGCCGCAACACAGGCTTCAGGCTTGTGAAGCAACCCGAGTTGGATGGACGAAAAG
ISM score
0.2
0.0
−0.2
Position
10
15
20
25
30
35
40
45
50
55
60
65
70
75
80
85
Position
0.0
0.1
0.2
0.3
0.4
0.5
0.6
Pairwise interaction score

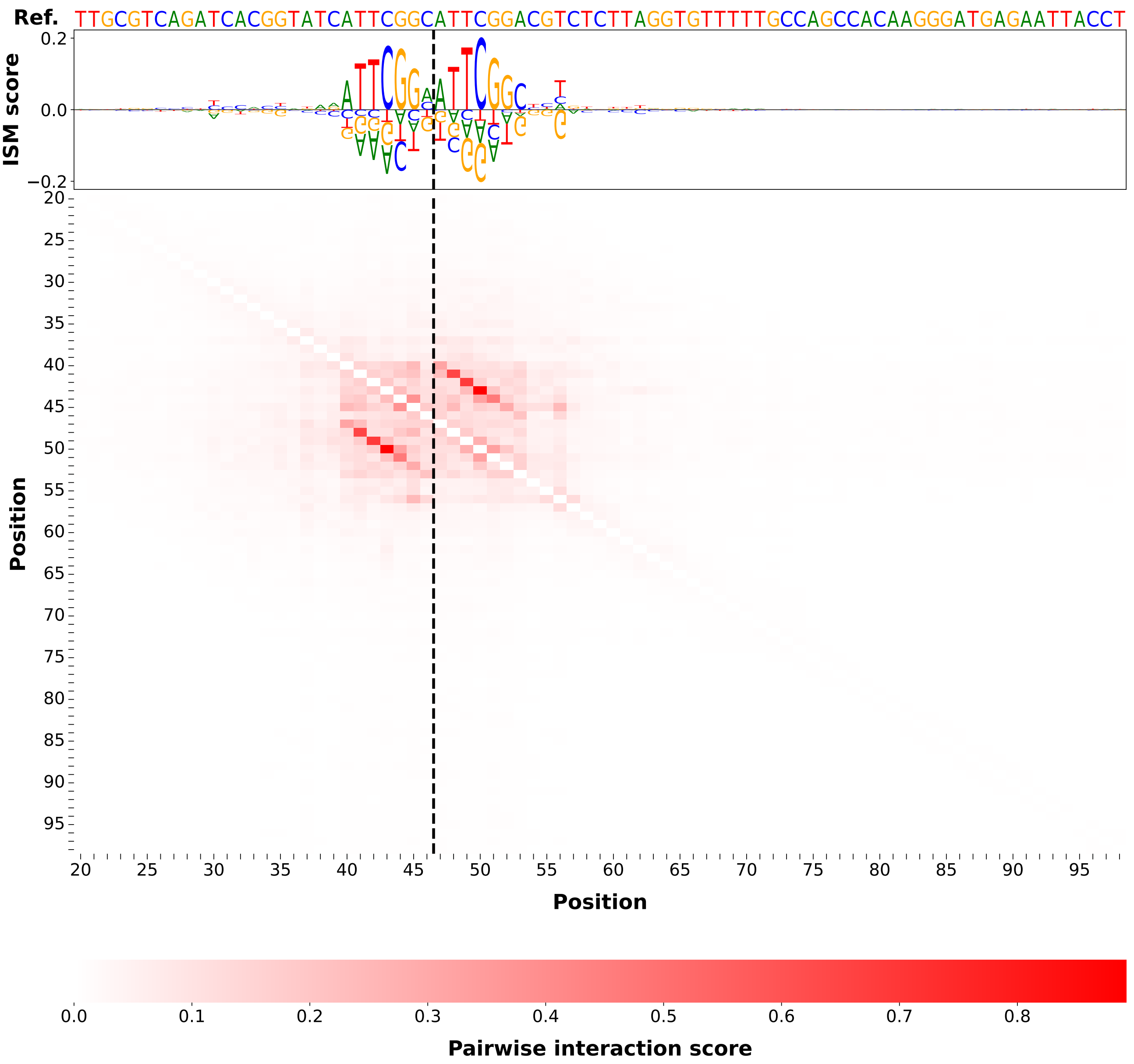
Ref. TTGCGTCAGATCACGGTATCATTCGGCATTCGGACGTCTCTTAGGTGTTTTTGCCAGCCACAAGGGATGAGAATTACCT
ISM score
0.2
0.0
−0.2
Position
20 25 30 35 40 45 50 55 60 65 70 75 80 85 90 95
Position
0.0 0.1 0.2 0.3 0.4 0.5 0.6 0.7 0.8
Pairwise interaction score

$\Delta$**length** $= -10$

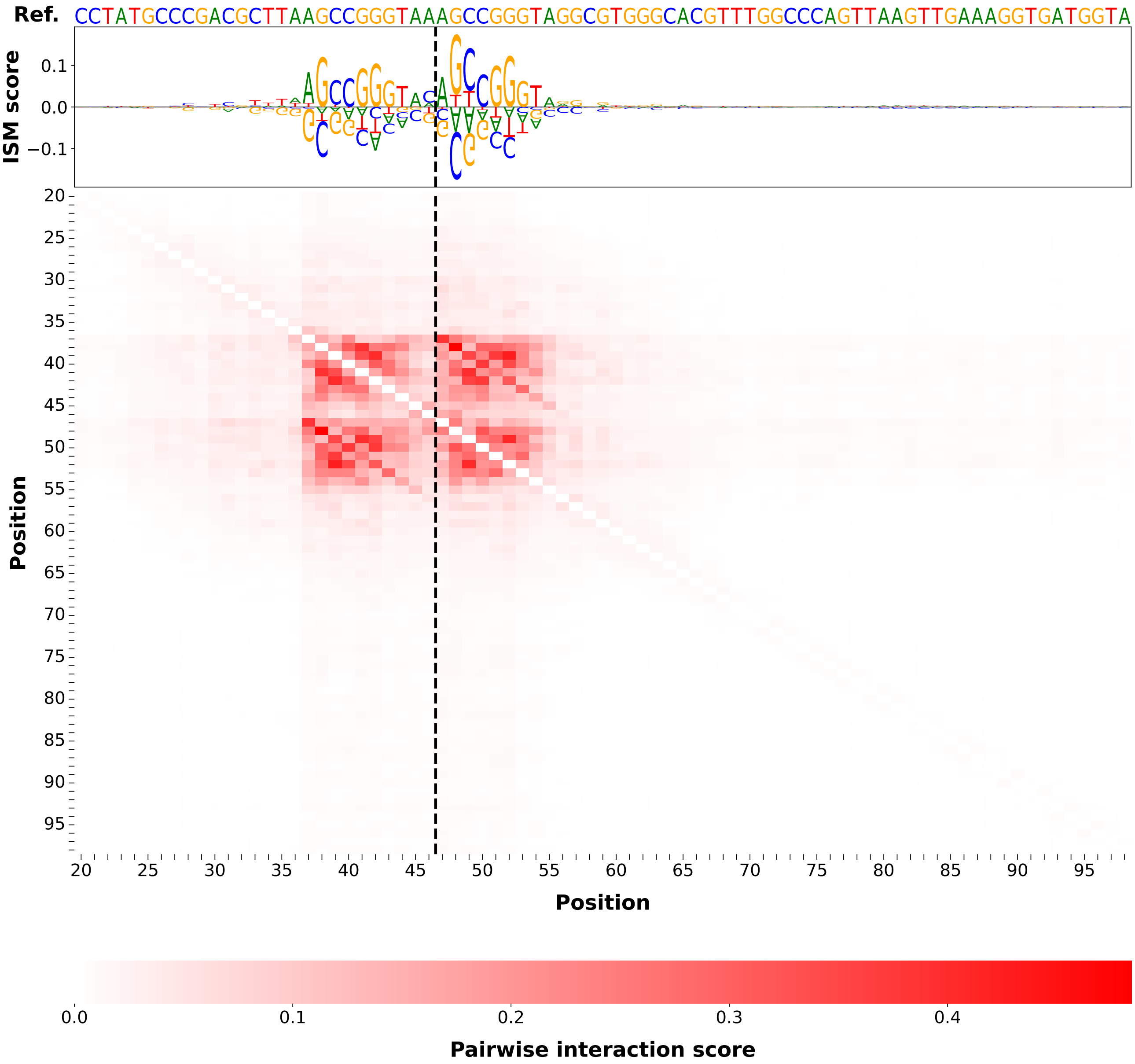

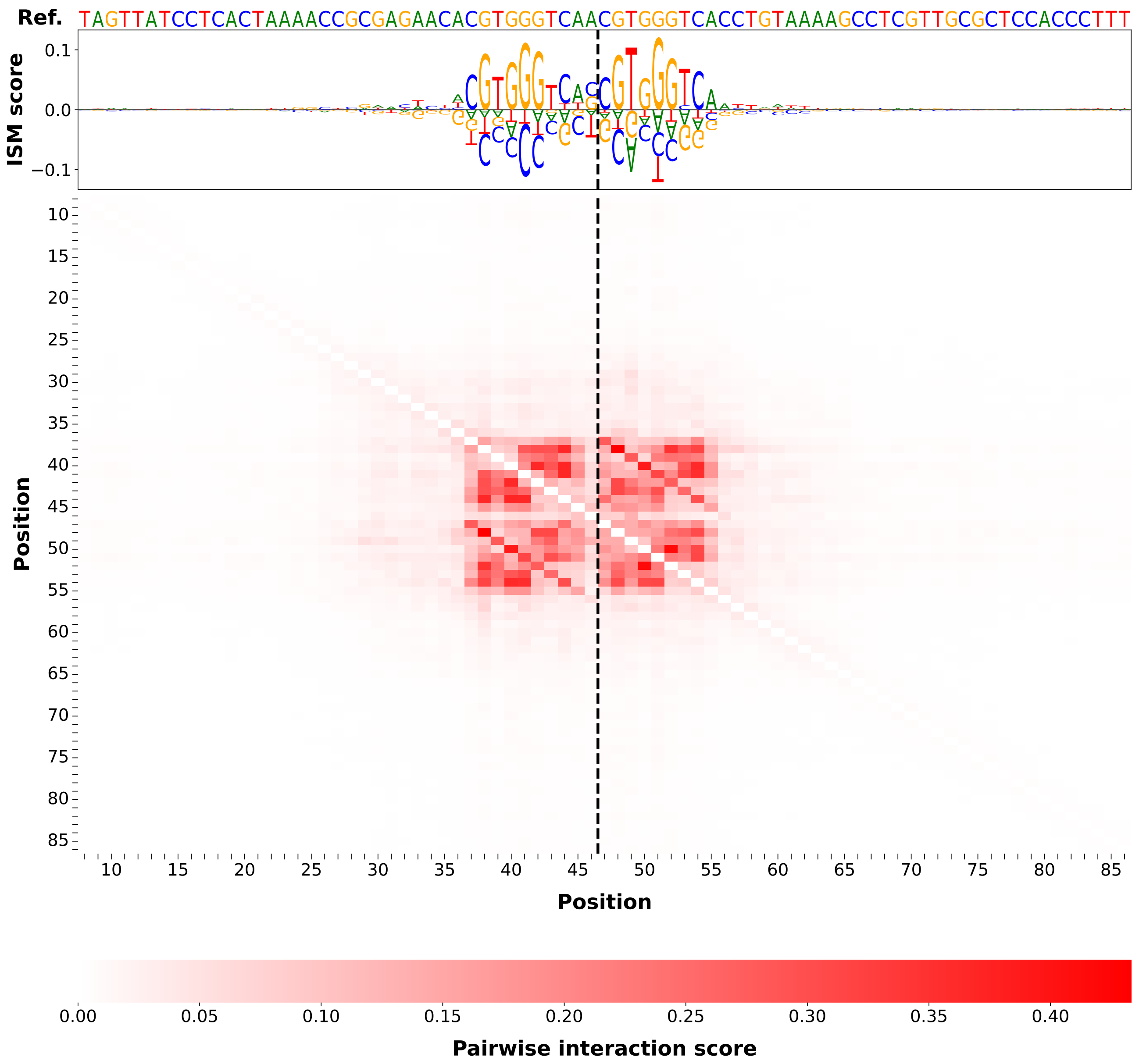
Ref. TAGTTATCCTCACTAAAACCGCGAGAACACGTGGGTCAACGTGGGTCACCTGTAAAAGCCTCGTTGCGCTCCACCCTTT
ISM score
0.1
0.0
−0.1
Position
10
15
20
25
30
35
40
45
50
55
60
65
70
75
80
85
Position
0.00
0.05
0.10
0.15
0.20
0.25
0.30
0.35
0.40
Pairwise interaction score

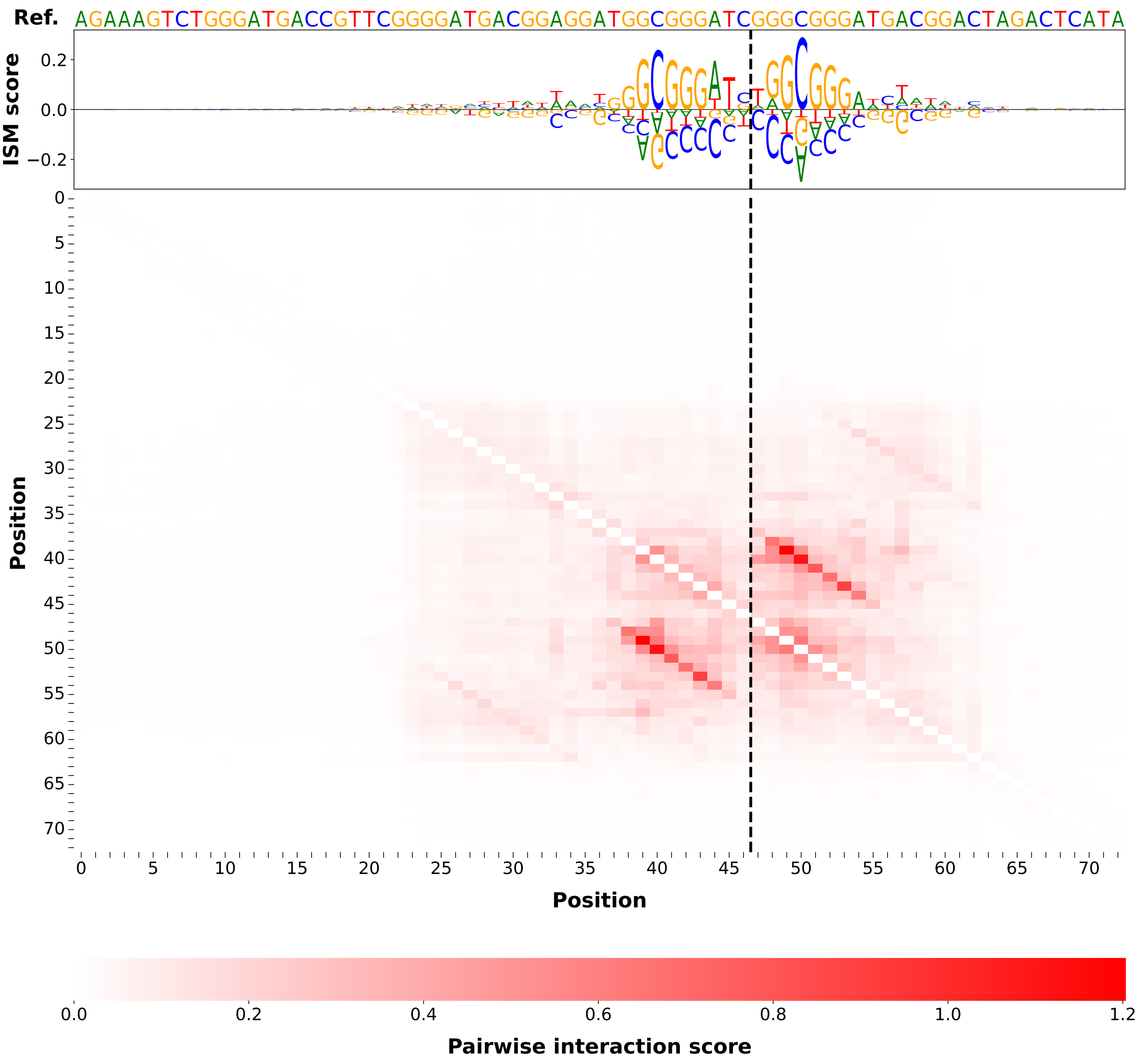
Ref. AGAAAGTCTGGGATGACCGTTCGGGGATGACGGAGGATGGCGGGATCGGGCGGGATGACGGACTAGACTCATA
ISM score
Position
Position
Pairwise interaction score

$\Delta$length $= -15$

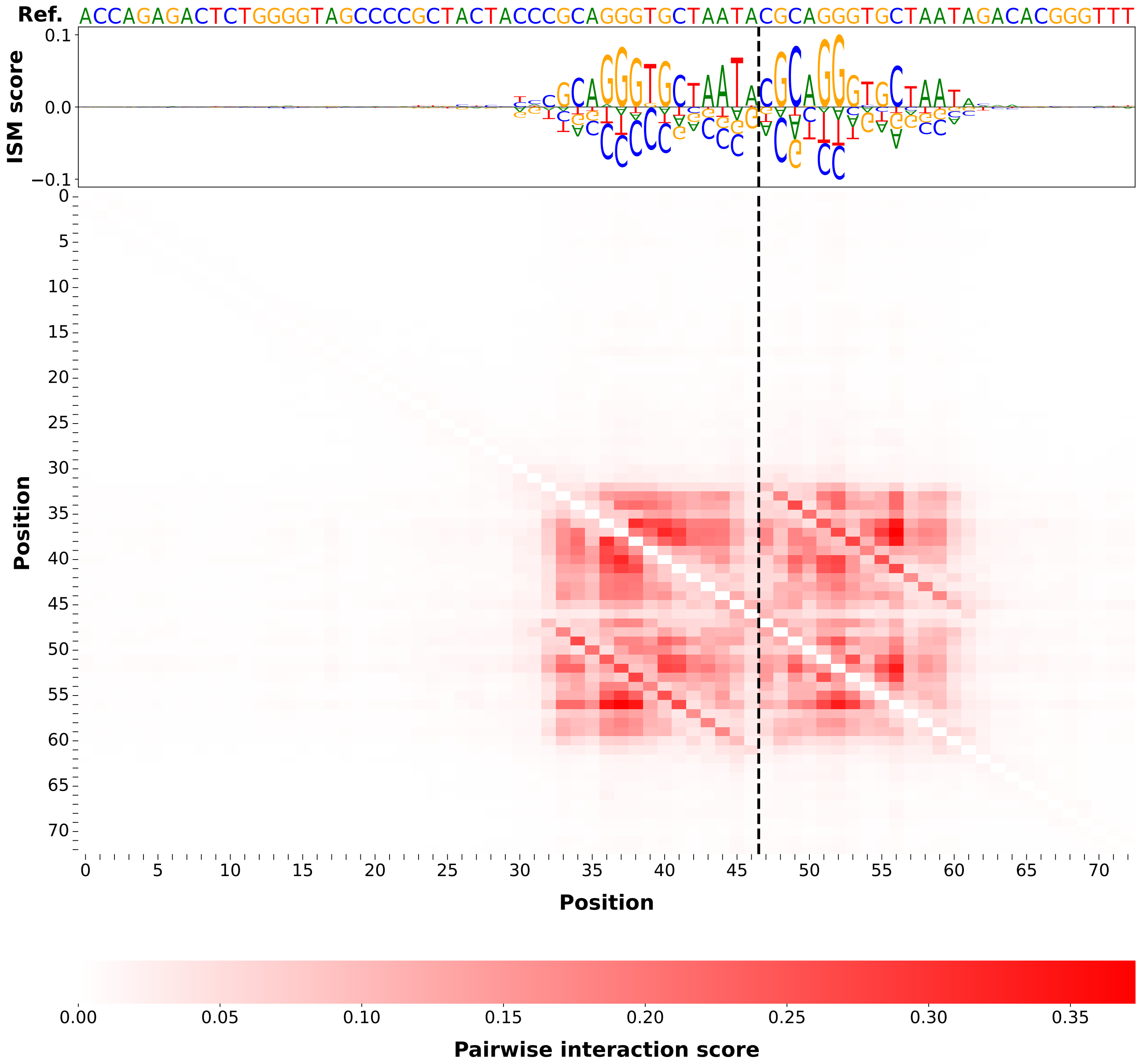

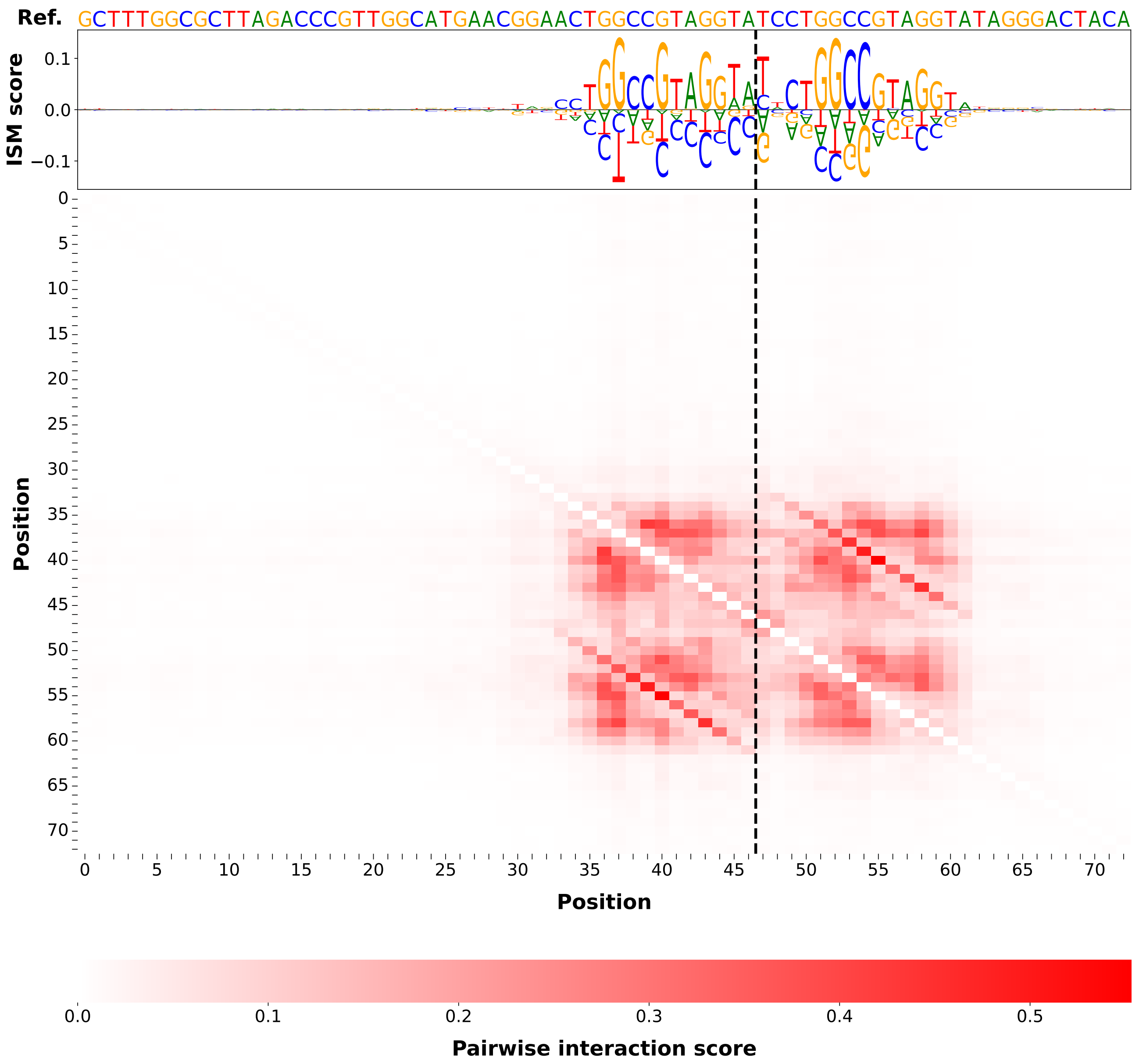
Ref. GCTTTGGCGCTTAGACCCGTTGGCATGAACGGAACTGGCCGTAGGTATCCTGGCCGTAGGTATAGGGACTACA
ISM score
Position
Position
Pairwise interaction score

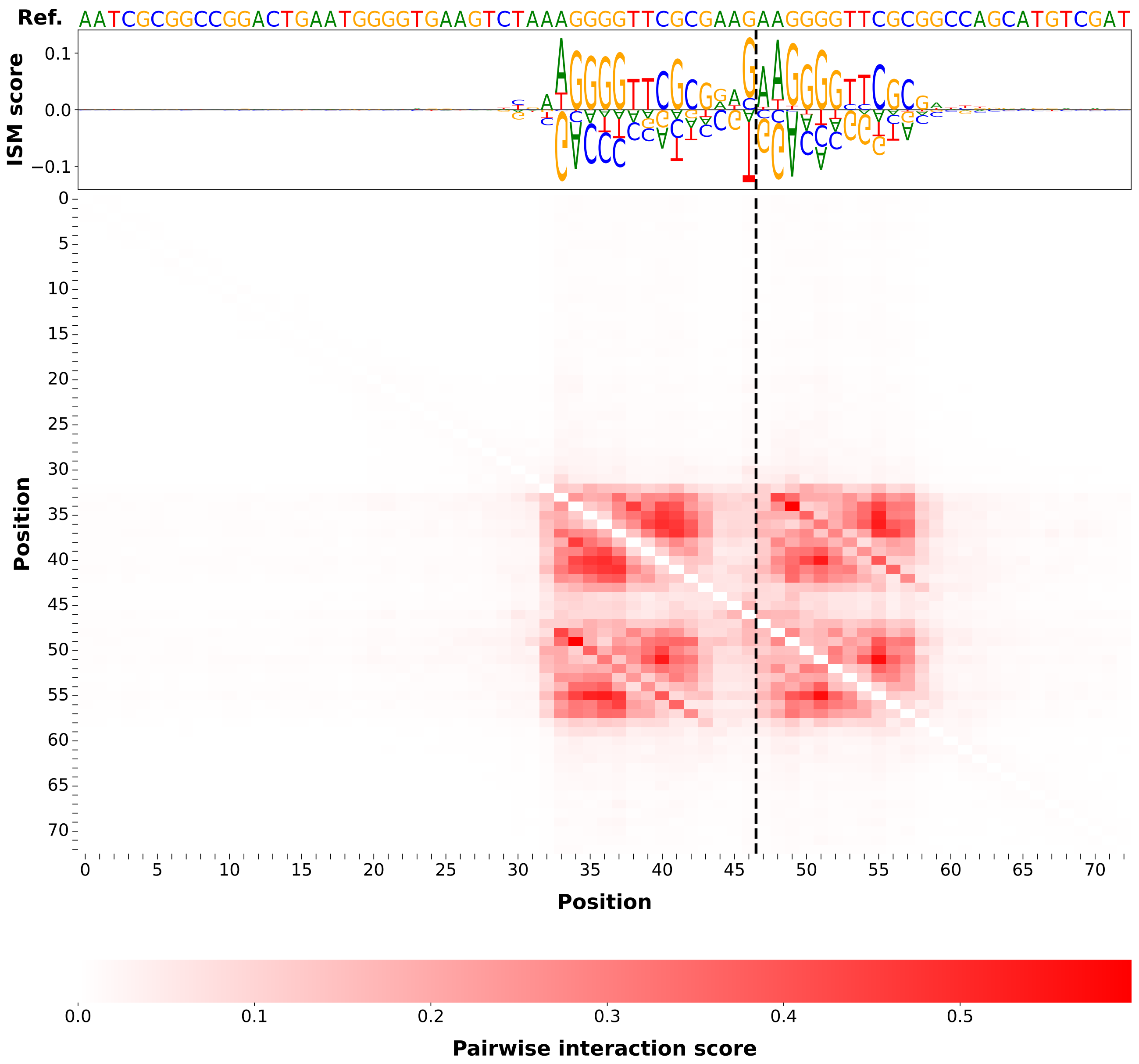
Ref. AATCGCGGCCGGACTGAATGGGGTGAAGTCTAAAGGGGTTCGCGAAGAAGGGGTTCGCGGCCAGCATGTCGAT
ISM score
0.1
0.0
−0.1
Position
0
5
10
15
20
25
30
35
40
45
50
55
60
65
70
Position
0.0
0.1
0.2
0.3
0.4
0.5
Pairwise interaction score

$\Delta$length $= -20$

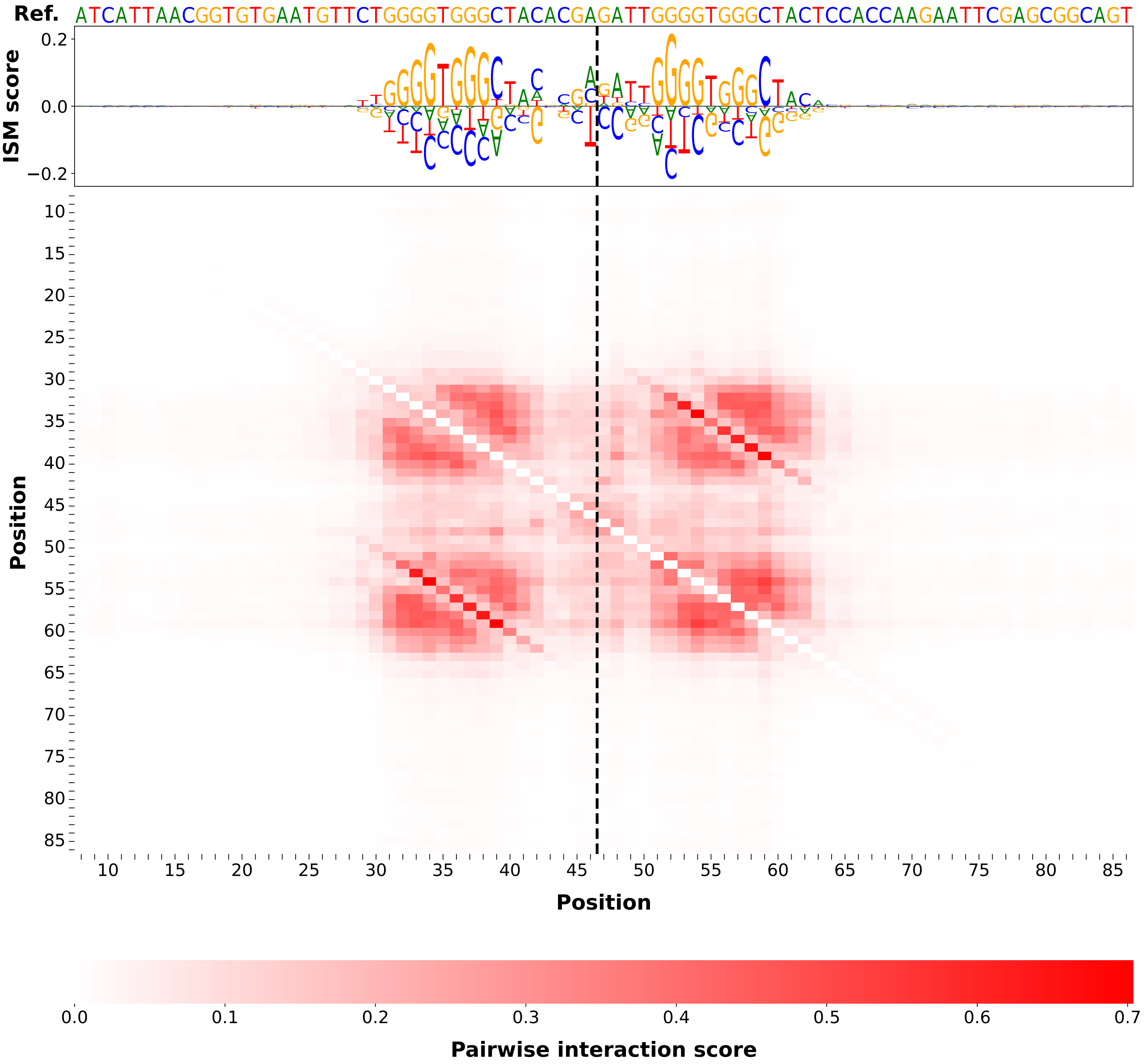

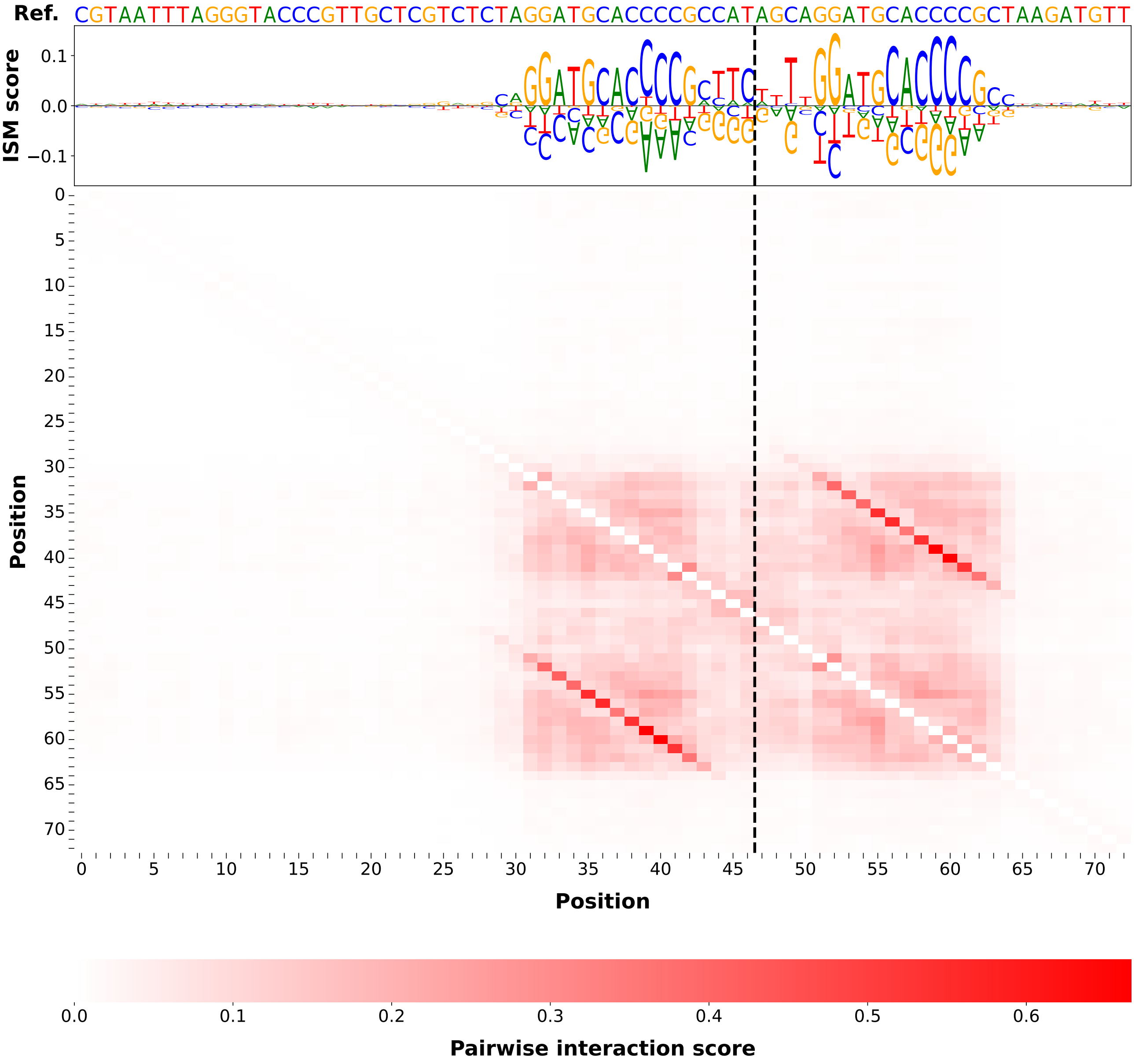
Ref. CGTAATTTAGGGTACCCGTTGCTCGTCTCTAGGATGCACCCCGCCATAGCAGGATGCACCCCGCTAAGATGTT
ISM score
Position
Position
Pairwise interaction score

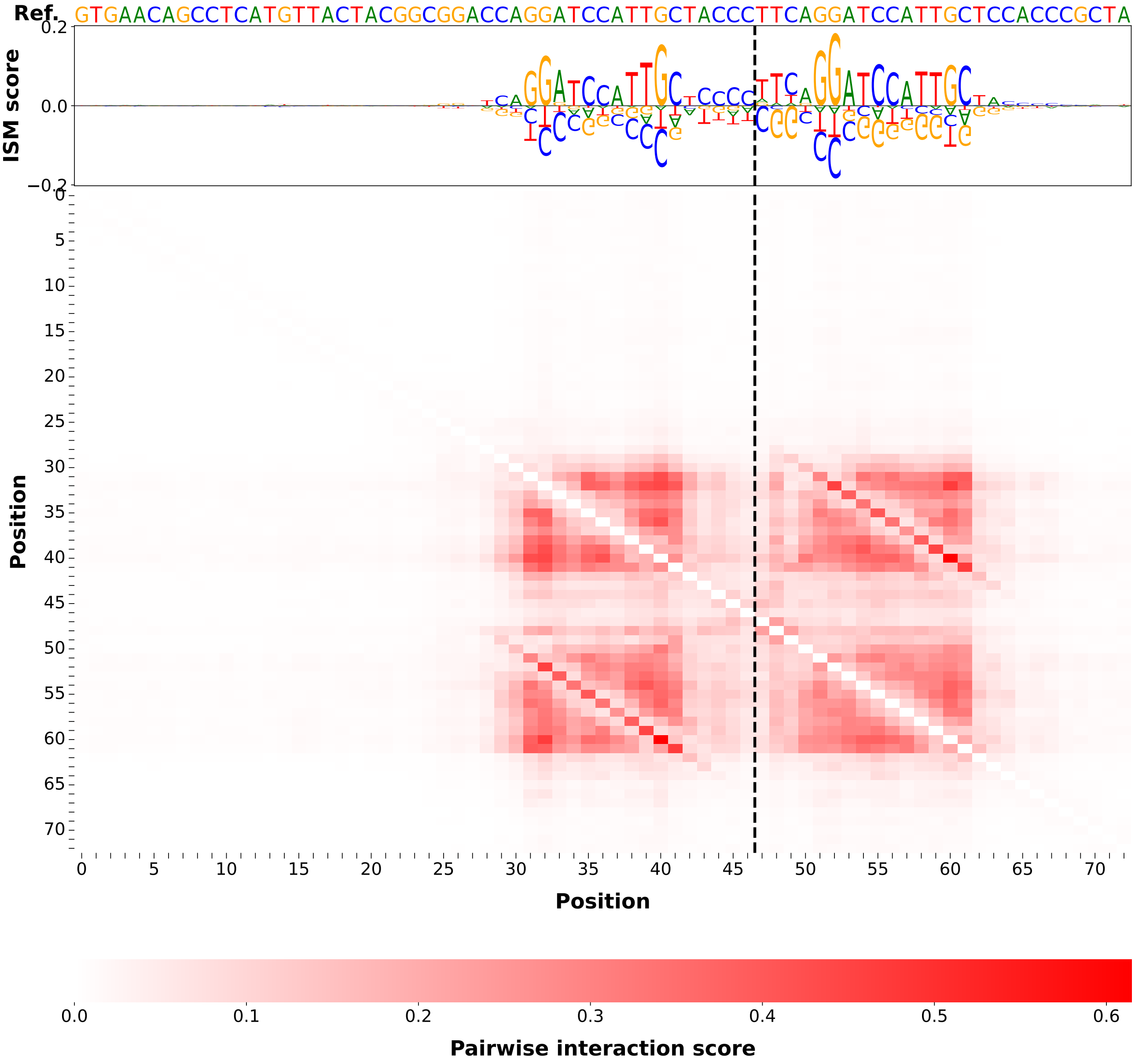
Ref.
GTGAACAGCCTCATGTTACTACGGCGGACCAGGATCCATTGCTACCCTTCAGGATCCATTGCTCCACCCGCTA
ISM score
Position
Position
Pairwise interaction score
